# Genetic diversity and population structure of *Epichloe* fungal pathogens of plants in natural ecosystems

**DOI:** 10.1101/2023.01.30.526252

**Authors:** Artemis D. Treindl, Jessica Stapley, Adrian Leuchtmann

## Abstract

Understanding the population genetic processes driving the evolution of plant pathogens is of central interest to plant pathologists and evolutionary biologists alike. However, most studies focus on host-pathogen associations in agricultural systems of high genetic and environmental homogeneity and less is known about the genetic structure of pathogen populations infecting wild plants in natural ecosystems. We performed parallel population sampling of two pathogenic *Epichloe* species occurring sympatrically on different host grasses in natural and seminatural grasslands in Europe: *E. typhina* infecting *Dactylis glomerata* and *E. clarkii* infecting *Holcus lanatus*. We sequenced 422 haploid isolates and generated genome-wide SNP datasets to investigate genetic diversity and population structure. In both species geographically separated populations formed genetically distinct groups, however population separation was less distinct in *E. typhina* compared to *E. clarkii*. The patterns of among population admixture also differed between species across the same geographic range: we found higher levels of population genetic differentiation and a stronger effect of isolation by distance in *E. clarkii* compared to *E. typhina*, consistent with lower levels of gene flow in the former. This pattern may be explained by the different dispersal abilities of the two pathogens and is expected to be influenced by the genetic structure of host populations. In addition, genetic diversity was higher in *E. typhina* populations compared to *E. clarkii*, indicative of higher effective population size in *E. typhina*. These results suggest that the effect of genetic drift and the efficacy of selection may differ in the two species. Our study provides evidence of how ecologically similar species occupying the same geographical space can experience different evolutionary contexts, which could influence local adaptation and coevolutionary dynamics of these fungal pathogens.

## 1 Introduction

Fungal plant pathogens are fundamental and ubiquitous components of natural biodiversity (Bradley *et al*. 2008; Burdon and Laine 2019). And yet, fungal plant pathogens are not commonly studied in natural ecosystems but rather in an agricultural setting, because they cause yield losses in important crop plants and may have negative impacts on human or animal health (Fisher *et al*. 2012). Evolutionary research in the field of plant pathology therefore heavily focuses on mechanisms leading to disease outbreaks in agricultural ecosystems usually associated with the rapid evolution of virulence or fungicide resistance related traits (McDonald and Linde 2002). Agricultural systems are useful for the study of antagonistic coevolution because these systems are characterized by strong selection and rapid adaptation, providing, in a way, short-term evolutionary “experiments” (Stukenbrock and McDonald 2008). A potential drawback, however, is the fact that the starting conditions of all these “experiments” are essentially the same: pathogens emerge under environmental and genetic uniformity. How plant pathogens evolve under more natural conditions remains less well understood. The impact of fungal plant pathogens on wild plants has received some attention in the case of invasive diseases [for example *Hymenoscyphus fraxineus* causing ash-dieback (McMullan *et al*. 2018)], or the chestnut blight fungus *Cryphonectria parasitica* (Milgroom *et al*. 2008), but mechanisms underlying persistence and evolutionary dynamics of host-pathogen associations in native systems remain much more elusive. Exploring the genetic structure of populations is central to the study of the evolutionary interactions between fungal pathogens and their host plants. The distribution of genetic variation within populations (genetic diversity) and among populations (genetic differentiation/patterns of gene flow) can provide insights into the evolutionary histories, distributions and disease dynamics of interacting species and determines their evolutionary potential (Thompson 2005, Chapter 2, Raw Materials for Coevolution I; Barrett *et al*. 2008). For example, the lack of genetic structure in host populations of cultivated plants favors the contagious spread of few (or a single) fungal pathogen genotypes and has been linked to devastating epidemics in agricultural systems (Stukenbrock and McDonald 2008; Persoons *et al*. 2017). In contrast, persistent antagonistic coevolution of pathogens with their hosts as it is observed in natural systems requires generation and maintenance of genetic variation in both partners. For example the analysis of natural populations of the flax rust pathogen *Melampsora lini* in Australia showed extensive genetic variation and population differentiation at loci associated with pathogenicity, suggesting that local evolutionary processes such as selection and genetic drift maintain high diversity between fungal pathogen populations (Barrett *et al*. 2009). In another well studied model pathosystem, the anther-smut fungus *Microbotryum lynchnidis-dioicae* infecting *Silene latifolia*, patterns of genetic structure were consistent with recolonization from ancient glacial refugia (Vercken *et al*. 2010; Badouin *et al*. 2017), and the population structure of the fungal pathogen was strongly congruent with the population structure of its host plant across the distribution range, a pattern likely driven by co-evolutionary processes (Feurtey *et al*. 2016). It appears that genetic structure of both the pathogen and the host underlies dynamics of disease prevalence at temporal and spatial scales and, for example in the case of powdery mildew *Podosphaera plantaginis* infecting *Plantago lanceolata*, this genetic and environmental heterogeneity among populations may buffer large-scale disease outbreaks (Jousimo *et al*. 2014). In order to increase our understanding of the evolutionary forces governing plant-pathogen interactions, more empirical studies on natural populations of fungal pathogens are needed that consider populations across a broad geographic range and encompass multiple comparable species.

Members of the ascomycete genus *Epichloe*, notably the *E. typhina* species complex, are an intriguing system in which to investigate the diversity and population structure of fungal pathogens in natural environments (Bultman et al. 2011). Fungi of the *E. typhina* complex reproduce sexually and sterilize their host plant while doing so. Since infections are systemic and long-lasting, the effect on host plant fitness is detrimental and infected plants may have no more reproductive output for the rest of their lifetime. In sterilizing systems such as this, interaction partners are engaged in a tight arms-race with strong selection pressures acting on the host plant to develop resistance and, on the pathogen, to overcome this resistance (Ashby and Gupta 2014; Tellier *et al*. 2014). While different *Epichloe* species are usually highly specialized on a single host grass species, these pathogens often occur in sympatric settings in natural grasslands hosting a variety of different grass species. This system thus provides an exceptional opportunity to sample sympatric populations and study ecologically similar pathogen species occupying the same geographic space.

In this study we sampled sympatric populations of two sibling species of the *E. typhina* complex occurring on distinct host grasses from natural grasslands across Europe. Using genome-wide SNPs we investigated population structure, that is the genetic diversity and patterns of differentiation within and among populations from different geographical regions. Our unique parallel sampling allows the analyses of both the genetic structure at the within-species level and a comparison of these patterns between the two species across the same geographic range. This provides important insights into how the biology of these species and the interaction with their host plants may have affected their evolutionary trajectories and helps to increase our understanding of the population genetic processes affecting fungal pathogens in natural ecosystems.

## 2 Materials and methods

### 2.1 Study system

#### 2.1.1 Fungal pathogens

The sexually reproducing members of the ascomycete genus *Epichloe* are highly specialized biotrophic endophytes and pathogens of many pooid grasses forming long-lasting systemic infections (Craven *et al*. 2001; Leuchtmann *et al*. 2014). Whole plants are usually infected by a single haploid genotype (Leuchtmann and Clay 1997), and obligate outcrossing takes place between two genotypes of opposite mating types, each infecting a different host individual (bipolar heterothallic mating system) (White and Bultman 1987; Schardl *et al*. 2014). As endophytes they form systemic and symptomless infections in vegetative tissues of the host grasses and then hijack grass flowering tillers to produce gametes and complete their own sexual cycle (Clay and Schardl 2002). The formation of the fungal fruiting structure (stroma) around the grass inflorescence causes choke disease, usually affecting all host inflorescences and therefore sterilizing the entire plant by preventing production of pollen and seed (Chung and Schardl 1997).

Here we focus on two closely related *Epichloe* species within the *E. typhina* species complex occurring on different Poaceae hosts: *E. typhina* infecting *Dactylis glomerata* and *E. clarkii* infecting *Holcus lanatus*. The two taxa are currently assigned the taxonomic rank of subspecies based on their sexual compatibility i.e. their ability to hybridize in experimental crosses (Leuchtmann and Schardl 1998; Leuchtmann *et al*. 2014). Indeed, hybrid ascospores between *E. typhina* and *E. clarkii* have been observed in natural sympatric populations (Bultman *et al*. 2011), indicating that at least the potential for gene-flow is retained. However, there is no evidence of host grasses infected by hybrid strains in natural populations suggesting the existence of postzygotic barriers. In line with our previous work which provided evidence of reproductive isolation and genome-wide high levels of genetic differentiation (Schirrmann *et al*. 2015, 2018; Treindl and Leuchtmann 2019), we consider *E. typhina* and *E. clarkii* as closely related yet distinct species at an advanced stage of divergence.

In a previous study, we generated high-quality genome assemblies of reference strains for both species which provide an essential genomic resource for the work conducted here (Treindl *et al*. 2021). Analyses of the structural organization of *Epichloe* genomes revealed a clear pattern of genome compartmentalization into gene-rich and AT-rich compartments, consistent with the so called “two-speed” model of genome evolution (see also Winter *et al*. 2018). This particular genome organization shared by some but not all plant-pathogenic fungi is thought to play an important role in driving rapid adaptation in the co-evolutionary arms-race between host and pathogen (Raffaele and Kamoun 2012; Möller and Stukenbrock 2017; Torres *et al*. 2020). A comparison of the *E. typhina* and *E. clarkii* reference genomes showed a high degree of genomic synteny between the sibling species, however, also revealed striking differences is genome size and compartmentalization likely related to an underlying difference in effective population size (Treindl *et al*. 2021). Besides the differences in genome structure and the obvious distinction based on host specificity, *E. typhina* and *E. clarkii* also differ in morphological characteristics including the size and disarticulation patterns of their ascospores (White Jr. 1993; Leuchtmann and Clay 1997).

#### 2.1.2 Host grasses

The host species *D. glomerata* (orchard grass or cocksfoot), and *H. lanatus* (Yorkshire fog) are both long-lived perennial grasses native to Europe (Tutin *et al*. 1980). They are wind-pollinated outcrossers, flowering typically May to July and are common and widely distributed in nutrient rich grasslands, where they often co-occur. Cultivars of *D. glomerata* may also be sown as part of commercial forage grass mixtures (e.g. see UFA-Samen, Herzogenbuchsee, Switzerland, and BARENBRUG France, Montévrain), while *H. lanatus* is not commercially sown. Although choke disease caused by *E. typhina* infections has been shown to cause yield reductions in commercial seed production of *D. glomerata* (Large 1954; Pfender and Alderman 2006), to our knowledge there has been no attempt to control pathogen prevalence in natural, semi-natural or pasture grasslands in Europe besides measures of selective breeding in commercial fields of seed producers and recommendations concerning mowing practices (Raynal 1991).

### 2.2 Sample collection and isolation of fungi

Sampling was performed at seven locations across Western Europe between May and July 2016-2018. At these locations large numbers (>20) of infected host grasses (*D. glomerata* and *H. lanatus* or, in one location *H. mollis*) occured intermixed or in close proximity less than 50 meters apart as evidenced by the presence of fungal fruiting bodies on flowering tillers (large circles in Figure 1, Table 1). The majority of the sampled populations were located in permanent natural meadows with no apparent evidence of recent high-intensity agricultural use such as regular sowing, fertilizing or grazing by livestock. One population (So) was located in a permanent semi-natural meadow which may be under low-intensity agricultural use. It is noteworthy that the transition between completely natural (no management) and semi-natural (extensive management) grasslands is fluid and even natural meadows often experience anthropogenic influences such as occasional mowing (e.g. in the Aub and Kew populations). However, *Epichloe* infected grasses were not observed in intensively managed meadows such as highly fertilized meadows or grass sown as an annual crop under crop rotation for high-intensity silage or hay production although single infected plants were observed around the rims of such fields (personal observation). It appears that agricultural practices that involve regular mowing (i.e. usually before host grasses flower) negatively affect the pathogen prevalence. We consider our populations to be «natural populations» in the sense that they are part of permanent grasslands not under strong human disturbance. From each host species 21-33 infected plants per location were sampled by picking one choked flowering tiller per plant individual. Each sample represents a different haploid genotype, since an individual plant is assumed to be infected by a single fungal strain resulting from horizontal transmission of meiotic ascospores (Leuchtmann and Clay 1997). In Kew Gardens (London, UK) *E. clarkii* was sampled from a previously unknown host species, *Holcus mollis*, occurring in sympatry with *E. typhina* on *D. glomerata. Holcus mollis*, unlike *H. lanatus*, can reproduce clonally through the formation of rhizomes in addition to normal sexual reproduction and prefers less nutrient rich sites in natural meadows or light woodland. (Ovington and Scurfield 1956). Therefore, we cannot exclude the possibility that some isolates from this population originate from a single fungal genotype systemically propagated via rhizomes within the population of *H. mollis*, although tillers were collected from plants at a distance of at least five meters. The sympatric population in Western Switzerland (Aub) had been sampled previously thirteen years prior to this study in 2005 (Steinebrunner and Leuchtmann, unpubl. data). We included these for sequencing and hereafter designate the isolates as Aub (sampled in 2018) and AubX (sampled in 2005). At locations where only one infected host species was found (small circles in Figure 1, Table 1) we collected infected flowering tillers from 1 to 4 individuals per population (Bret, Other 1-4) and refer to these as “additional” samples.

**FIGURE 1.**
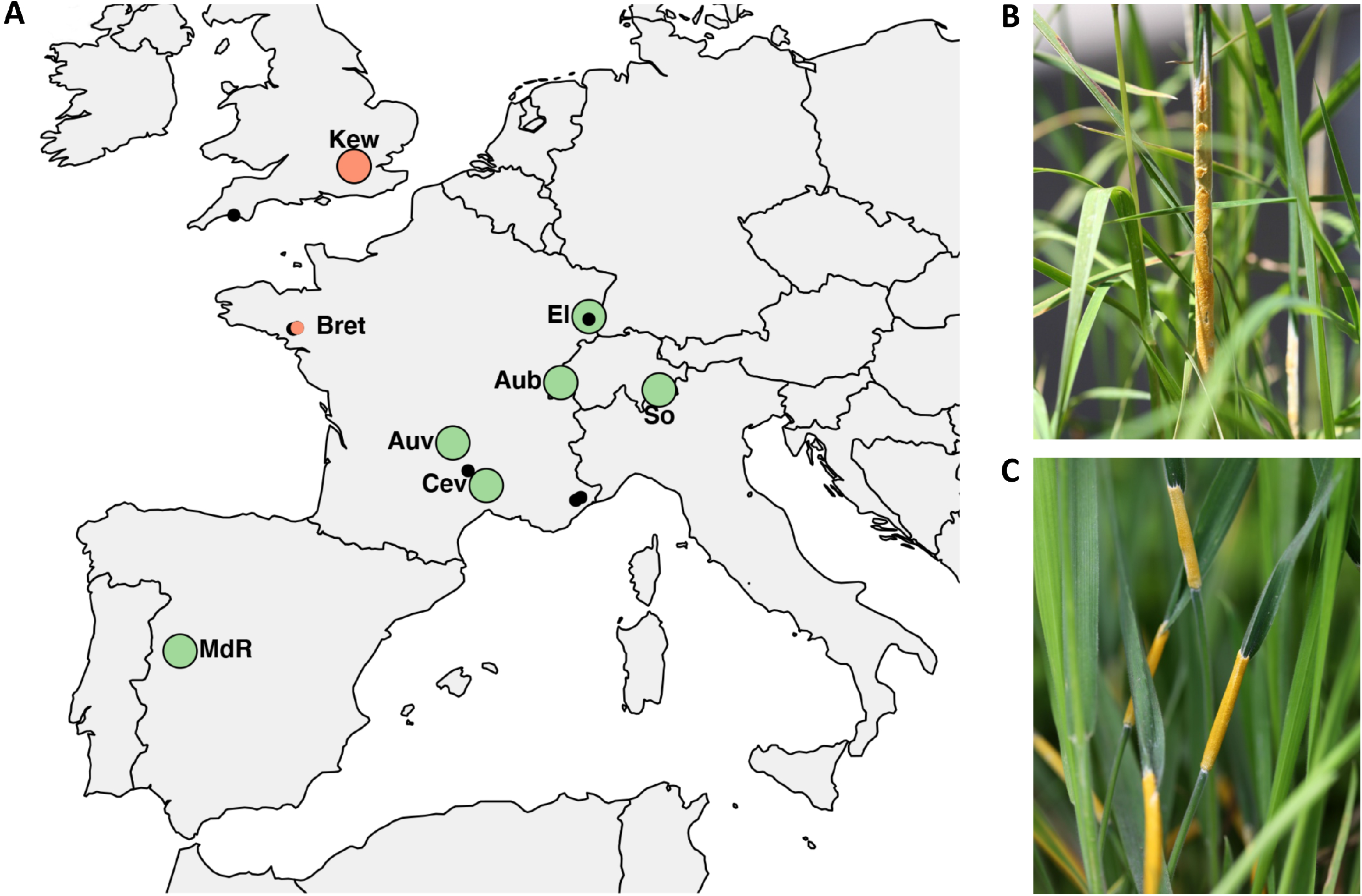
(**A**) Populations of *Epichloe* isolates across Europe used in the study. Large circles indicate sampling locations of sympatric populations where *E. typhina* is infecting *D. glomerata* and *E. clarkii* is infecting *H. lanatus* (green) or *H. mollis* (red), respectively. Smaller circles indicate additional samples collected from allopatric populations where only one of the two host species was found to be infected, either *H. mollis* (red) or *D. glomerata* (black). Among these are five locations (small black circles) where *E. typhina* occured alone (n=2-4), collectively designated as “Other”. The population in Western Switzerland (Aub) was sampled twice thirteen years apart, in 2005 and 2018. (**B**) Ripe fungal fruiting body (stroma) of *E. typhina* infecting *D. glomerata*. (**C**) Stromata of *E. clarkii* infecting *H. lanatus*. For code and origin of populations see Table 1.

**TABLE 1.** Populations of *Epichloe* infecting *Dactylis* and *Holcus* host species with sampling location, year of collection, host and fungal species, and number of isolates (n) analyzed in this study.

| Population ID | Sampling location | Latitude | Longitude | Collection year | Host species | Fungal species | n |
| --- | --- | --- | --- | --- | --- | --- | --- |
| Aub | Aubonne, VD, Switzerland | 46.5123 | 6.3637 | 2018 | *D. glomerata* | *E. typhina* | 22 |
|  |  |  |  |  | *H. lanatus* | *E. clarkii* | 32 |
| AubX | Aubonne, VD, Switzerland | 46.5123 | 6.3637 | 2005 | *D. glomerata* | *E. typhina* | 23 |
|  |  |  |  |  | *H. lanatus* | *E. clarkii* | 23 |
| Auv | Murat, Auvergne, France | 45.1263 | 2.8863 | 2017 | *D. glomerata* | *E. typhina* | 30 |
|  |  |  |  |  | *H. lanatus* | *E. clarkii* | 32 |
| Cev | Saint-Paul-la-Coste, Cevennes, France | 44.1483 | 3.9627 | 2017 | *D. glomerata* | *E. typhina* | 31 |
|  |  |  |  |  | *H. lanatus* | *E. clarkii* | 31 |
| El | Abbaye de Marbach, Alsace, France | 48.0266 | 7.2752 | 2017 | *D. glomerata* | *E. typhina* | 32 |
|  |  |  |  |  | *H. lanatus* | *E. clarkii* | 32 |
| Kew | Kew Gardens, London, UK | 51.4783 | -0.2990 | 2017 | *D. glomerata* | *E. typhina* | 21 |
|  |  |  |  |  | *H. mollis* | *E. clarkii* | 30 |
| MdR | Montemayor del Rio, Spain | 40.3469 | -5.9091 | 2018 | *D. glomerata* | *E. typhina* | 33 |
|  |  |  |  |  | *H. lanatus* | *E. clarkii* | 29 |
| So | Soglio, GR, Switzerland | 46.3429 | 9.5373 | 2016 | *D. glomerata* | *E. typhina* | 27 |
|  |  |  |  |  | *H. lanatus* | *E. clarkii* | 33 |
| Bret<sup>a</sup> | La Gacilly, Bretagne, France | 47.7692 | -2.1384 | 2017 | *D. glomerata* | *E. typhina* | 4 |
|  |  | 47.7687 | -2.1300 | 2017 | *H. mollis* | *E. clarkii* | 3 |
| Other 1<sup>a</sup> | Mount Edgecomb, Plymouth, UK | 50.3554 | -4.1753 | 2017 | *D. glomerata* | *E. typhina* | 3 |
| Other 2<sup>a</sup> | Cultures, Mende, France | 44.4890 | 3.3751 | 2017 | *D. glomerata* | *E. typhina* | 3 |
| Other 3<sup>a</sup> | Haut Thorenc, Préalpes d'Azur, France | 43.8743 | 7.0097 | 2017 | *D. glomerata* | *E. typhina* | 6 |
| Other 4<sup>a</sup> | Westhalten, Alsace, France | 47.9656 | 7.2696 | 2017 | *D. glomerata* | *E. typhina* | 4 |

<sup>a</sup> Additional samples from allopatric populations

Fungal strains were isolated from freshly collected tillers by splitting open the stromata with a sterile blade to expose the interstitial fungal mycelium between undeveloped inflorescence and leaf blades. Pieces of mycelium were then transferred with sterile tweezers and subsequently grown on supplemented malt extract-agar (SMA) plates containing 1 % malt extract, 1 % glucose, 0.25 % bacto peptone, 0.25 % yeast extract, 1.5 % bacto agar and 0.005% oxytetracycline (Pfizer, New York, NY, U.S.A.). Isolates were carefully checked under the dissecting scope for purity and *Epichloe* identity as described previously (Christensen *et al*. 1993). Pure cultures were grown in V8 liquid media on a rotary shaker at 12 rpm for 10-14 days at room temperature. Mycelium was vacuum filtered, freeze dried for 24 h and then ground using liquid nitrogen. All live axenic cultures were also stored on SMA in tubes covered with mineral oil at 4 °C at ETH Zurich.

### 2.3 DNA preparation and sequencing

Genomic DNA was extracted from 10 mg freeze dried material using the sbeadex mini plant kit (LGC Genomics, Berlin, Germany), but replacing the lysis buffer with PVP (sbeadex livestock kit) as this buffer had previously shown improved DNA yields. Extraction was automated on a KingFisher system following the manufacturers protocol (Thermo Fisher Scientific, Waltham, MA, USA). The DNA concentration of extracted samples was quantified using a Spark^®^ Multimode Microplate Reader (Tecan Trading AG, Switzerland). Paired-end libraries of 2 x 150 bp fragments with an insert size of 330 bp were prepared with NEBNext^®^ Ultra™ II DNA Library Prep Kits (New England Biolabs, Ipswich, MA, USA), and sequencing was performed on a HiSeq4000 Illumina sequencer (Illumina, San Diego, CA, USA), at 16x coverage on average. We sequenced whole genomes of 240 individuals of *E. typhina* and 240 individuals of *E. clarkii* of which 211 each yielded sufficient data for the analyses. All Illumina sequence data will be made accessible on the NCBI Short Read Archive upon publication.

### 2.4 Mapping and variant calling

Reads were mapped against the high-quality reference genomes of *E. typhina* (Ety_1756, BioProject ID PRJNA533210) and *E. clarkii* (Ecl_1605_22, BioProject ID PRJNA533212) (Treindl *et al*. 2021). Mapping was performed with BWA (Li and Durbin 2009) and variant calling was done with the Genome Analysis Toolkit (McKenna *et al*. 2010), using the GATK Short Variant Discovery best practice workflow (https://gatk.broadinstitute.org/hc/en-us/articles/360035535932-Germline-short-variant-discovery-SNPs-Indels-). Details and examples of how each command was run are provided on DRYAD (for review purpose these are hosted temporarily on github (https://github.com/adtreindl/Epichloe_genomics). In brief, the raw reads were trimmed and adaptors removed using trimommatic v0.35 (Bolger *et al*. 2014). The trimmed reads were mapped to the respective reference genomes using bwa mem v0.7.17 (Li and Durbin 2009). Low quality alignments (qual>10) were removed and duplicates were marked. Mapping statistics were obtained using sambaba v0.6.6 (Tarasov *et al*. 2015). After mapping, variants were identified using HaplotypeCaller, GatherVCFs and JointGenotyper.

### 2.5 Filtering

Variants were filtered with custom scripts using vcftools (Danecek *et al*. 2011), bcftools (Li *et al*. 2009), and R (R Core Team 2013) (for example scripts see github repository (https://github.com/adtreindl/Epichloe_genomics). We removed mitochondrial sequences and indels, thus only SNPs in nuclear DNA (Chr1-7) were used in all subsequent analysis. The following hard-filters were applied using vcftools v1.2.8 minor allele count (--mac 3) and maximum mean depth values (--max-meanDP 100). The latter filter removed variants with excessive coverage (mean depth >> mean read depth) across all samples as these are indicative of either repetitive regions or paralogs. We filtered out SNPs that were not well sampled across populations by removing SNPs with >30% missing within a single population and removed isolates that had >80% missing data. We then filtered by read quality and read depth following GATK best practice recommendations (Van der Auwera *et al*. 2013), and based on our evaluation of the data. We removed sites with low mapping quality (MQ value < 40, the Root Mean Square of the mapping quality of the reads across all samples) and with a high relative variance in read depth (variance in read depth divided by mean read depth > 80 for *E. typhina* and > 30 in *E. clarkii* (Supplementary Figure S1). We visually inspected regions at different SNP densities using Integrative Genomics Viewer (IGV(Robinson *et al*. 2011) to assess the variant calls and based on these observations chose to excluded sites where ≥3 SNPs had been called within 10bp. Lastly, we removed sites that previous filtering steps had rendered monomorphic. Summary statistics including missingness and mean depth of this dataset are shown in Supplementary Figure S2. For some downstream analyses additional filters were applied and are described below and listed in Supplementary Table S1.

### 2.6 Analyses of genetic structure

We analyzed genetic structure of 211 *E. typhina* and 211 *E. clarkii* isolates using several methods: Principal component analysis (PCA), Bayesian clustering analysis (STRUCTURE) and discriminant analyses of principal component (DAPC). PCA, which does not rely on any population genetic model, was implemented in the R package SNPRelate (Zheng *et al*. 2012). PCAs were done for the complete SNP datasets, LD-pruned datasets (see below) and datasets with no missing data. We repeated analyses after removing the MdR population from the datasets, as these individuals are so genetically distinct in both species that they obscure the genetic structure of the other populations. We also visualized the amount of sub-structure within each population using PCA (see Supplementary Notes: Population sub-structure). For the STRUCTURE analysis, we applied a minor allele frequency filter of 0.05 using vcftools (–maf 0.05) to exclude rare alleles from the analysis as they are likely not representative of a particular population group. We also removed SNPs that are in linkage disequilibrium (LD), pruning variants with a pairwise r^2^ greater than 0.3 in 10kb windows using bcftools v1.9. The LD-pruned datasets included 152,883 biallelic SNPs in *E. typhina* and 79,352 SNPs in *E. clarkii*. STRUCTURE v2.3.4 (Pritchard *et al*. 2000), was run for K=1-12 with 20,000 Monte Carlo Markov Chain (MCMC) iterations following a 10,000 iteration burn-in for each of twelve replicate runs per K. We used an admixture ancestry model with correlated allele frequencies and no prior information about the demography. The output of the STRUCTURE analysis was processed using STRUCTURE HARVESTER (Earl and vonHoldt 2012). The model with the maximized rate of change in log likelihood values (ΔK) was considered the optimal number of clusters based on methods described by Evanno et al. (Evanno *et al*. 2005). We assessed probability of assignment and genetic admixture among populations by plotting the proportion of membership to each of the inferred clusters for every individual, using the optimal model parameters (the best run) for each K.

DAPC was performed using the R package *adegenet* v2.0.0 (Jombart 2008) and the ‘find.clusters’ function. This method first transforms the data using PCA and then performs a Discriminant Analysis on the retained principal components. For computational reasons we used the LD-pruned dataset and further reduced the number of SNPs to one variant per 2kb using vcftools (--thin 2000). This left 12,562 SNPs in *E. typhina* and 11,338 SNPs in *E. clarkii* (LD-pruned and thinned dataset).

### 2.7 Summary statistics of genetic variation

Population genetic summary statistics were calculated with vcftools v1.16, using a patch that allows computation with haploid datasets (https://github.com/vcftools/vcftools/pull/69). Calculations were performed for the seven sympatric populations only (additional samples from allopatric locations were removed), leaving 191 individuals in *E. typhina* and 208 individuals in *E. clarkii*. After removing “additional” isolates we again removed any monomorphic variants. Populations sampled in different years from the Aubonne location (Aub and AubX) were analyzed as separate populations. Step-by-step details and example code is available at: https://github.com/adtreindl/Epichloe_genomics/blob/master/summary_stats.md

#### 2.7.1 Nucleotide diversity π, Tajimas D

We quantified within-population genetic diversity by calculating nucleotide diversity in 10kb non-overlapping windows (π, the average number of differences between individuals (Takahata and Nei 1985)), and Tajima’s D in 10kb and 40kb non-overlapping windows (Tajima 1989). Tajima’s D reflects the distribution of allele frequencies within populations also called the site frequency spectrum. Tajima’s D is > 0 when there is a lack of rare alleles indicative of balancing selection or population contraction, Tajima’s D < 0 indicates an excess of low frequency alleles as a result of recent positive selection or population expansion, and Tajima’s D of 0 suggest neutral evolution. Genomic regions with reduced levels of nucleotide diversity and Tajima’s D may indicate departure from neutrality that can be the result of demographic processes or positive selection acting on these loci. We calculated the average nucleotide diversity and Tajima’s D per population as the mean across all windows.

#### 2.7.2 F_ST_

To assess relative genetic differentiation between populations we calculated pairwise fixation indexes FST (Weir and Cockerham 1984), per site between all populations using vcftools (haploid switch, see above). We then calculated mean F_ST_ in non-overlapping windows of 5kb (10kb) along each chromosome using R version 4.0.2 (R Core Team 2013), and genome-wide weighted mean F_ST_ between population pairs across all SNPs. To investigate patterns of isolation-by-distance (IBD) in *E. typhina* and *E. clarkii* we tested for a positive correlation between genetic distance (the mean weighted FST) and geographic distance. We used the R package lme4 (Bates *et al*. 2015) to fit a linear mixed-effects model to the data with populations as random effects and distance as a fixed effect. We also calculated divergence as the number of fixed differences between population pairs (SNPs with an F_ST_ of 1).

#### 2.7.3 Analysis of linkage disequilibrium

We removed loci with a minor allele frequency below 0.05 (within a population) because differences in allele frequencies can (upwardly) bias estimation of LD. We calculated the coefficient of correlation (r^2^) between pairs of SNPs that were up to 20 kb apart using plink (v. 1.90) and calculated mean LD within 100bp windows. We visualized the decay of r^2^ over physical distance using a locally weighed polynomial regression (LOESS) model using the function *geom_smooth* in the R package ggplot2 (Wickham 2016).

#### 2.7.4 Effective population size N_e_

To compare effective population sizes within and between species we estimated effective population sizes (*N*_e_) from polymorphism data with the parameter θ = 2*N*_e_μ (for more details see Supplementary Notes: Calculation of effective population size). Calculations were based on Watterson’s estimators of θ (Watterson 1975), which is based on the number of segregating sites in a population relative to the sequence length, and an assumed mutation rate (μ) of 3.3 × 10^-8^ per site. This mutation rate is based on estimates in yeast (Lynch *et al*. 2008) and has been used in other studies (Stukenbrock *et al*. 2011). The number of segregating sites per populations can be found in the Supplementary (Table S2). From this we also calculated the proportion of segregating sites within each population relative to the total variation among all populations, as a measure of genetic diversity comparable between species (Supplementary Table S3).

We also analyzed the distribution of genetic variation separately for distinct genomic compartments in the *Epichloe* genomes. Methods and results are presented in the Supplementary (Supplementary Notes: PCA analysis and calculation of summary statistics by compartments). Additionally, we assessed population sex-ratios (Supplementary Notes: Mating types) and interspecific hybridization in two populations (Supplementary Notes: Assessment of hybridization in sympatric populations).

## 3 Results

### 3.1 Genetic structure of populations of two *Epichloe* species

After removal of isolates with low coverage and replicates we retained genome-wide polymorphic sites for 211 isolates in each of the species. We obtained 658,021 biallelic SNPs in the *E. typhina* dataset and 400,033 SNPs in the *E. clarkii* dataset after initial filtering (complete dataset, Supplementary Table S1).

Both Principal component (PC) and Bayesian clustering found that population structure of the two species reflected the geographic origin of sampled populations, however, patterns of genetic differentiation differed between species and STRUCTURE indicated fewer clusters in *E. typhina* with three populations (Kew, Auv, El) being assigned to the same cluster (see below). PCA grouped genotypes according to their sampling location, with the Spanish population (MdR) forming the most genetically distinct group (Supplementary Figure S3A and D). We removed this population in both species and repeated the analysis to explore genetic patterns among the remaining populations (Figure 2 and Figure 3). Clusters representing the sampled *E. typhina* populations could be clearly distinguished in the principal component space with the first axis resolving the West-to-East distribution (explaining 3.93% of the variation), and the second axis resolving the South-to-North distribution (2.58% variation explained, Figure 2B). Spatially intermediate “additional” samples (Bret and Other 1-4) were also genetically intermediate. In *E. clarkii* seven clusters were evident in the PCA, each corresponding to a single geographic population with the exception of the Kew population, which consisted of two genetically distinct groups (Figure 3B). In both species, isolates sampled thirteen years apart from the same Western Switzerland location did not form genetically distinct groups (AubX and Aub sampled 2005 and 2018), suggesting little genetic differentiation has occurred during this time. These population genetic structure results did not differ between datasets filtered differently as outlined in the methods section (complete, LD pruned, no missing data) (Supplementary Figures S3 and S4). The two populations that were the most spread out in principal component space in overall analyses (*E. typhina* Cev and *E. clarkii* Kew) also showed evidence of sub-structure in PCA performed within populations using only isolates from the same location (see Supplementary Notes and Supplementary Figures S5 and S6).

**FIGURE 2.**
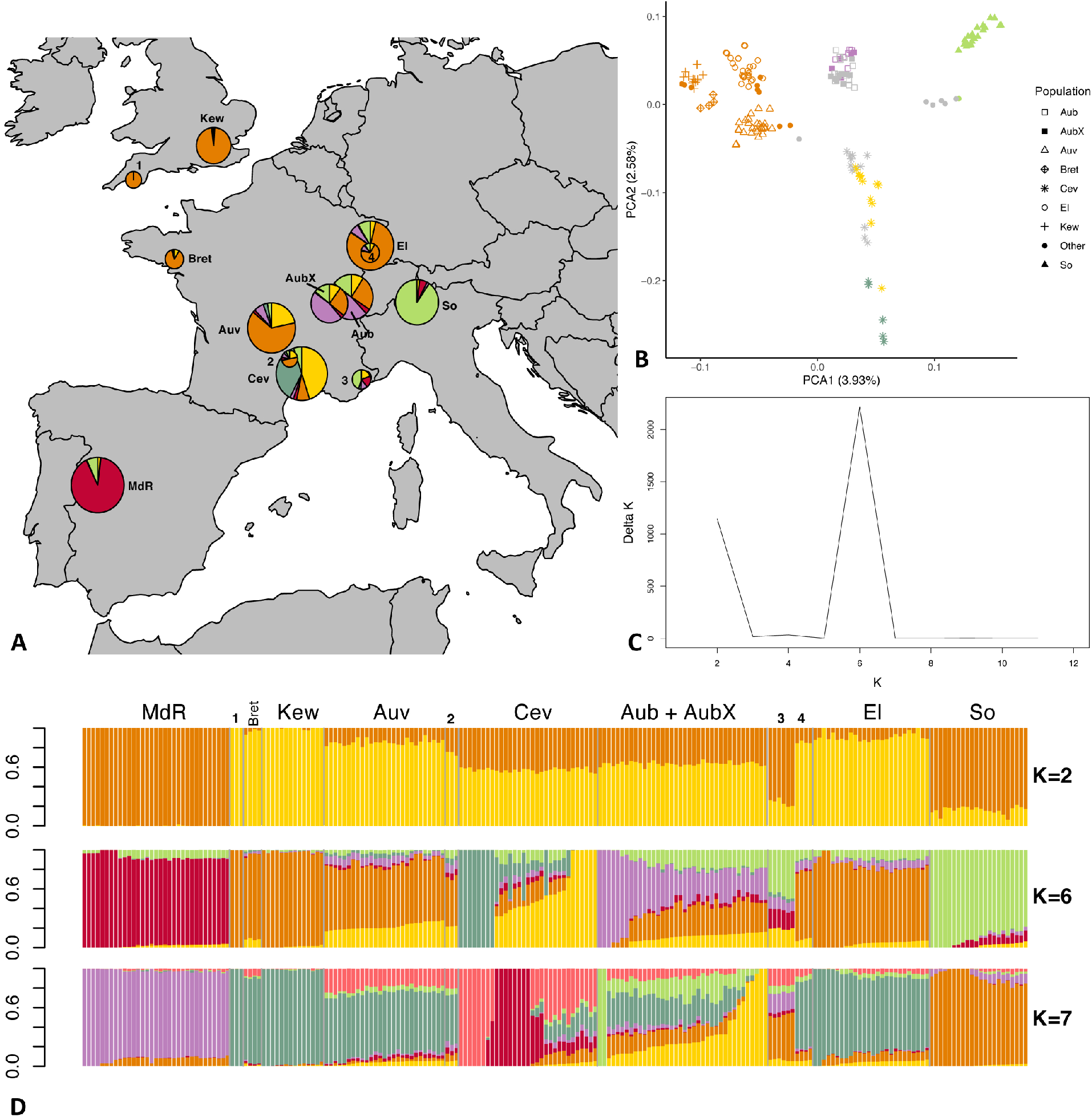
Population structure analyses of 211 *E. typhina* isolates based on genomewide SNPs. The LD pruned dataset including 152,883 SNPs was used in the STRUCTURE analyses whereas the complete dataset of 658,021 SNPs was used for the PCA. (**A**) Sampling locations of *E. typhina* isolates infecting *D. glomerata*; pie charts represent Bayesian cluster membership proportion for K=6, diameters reflect sample size (range n=3 to n=33). (**B**) Principal component analysis of genetic differentiation among isolates excluding the population from Spain (MdR); the percentage of variance explained by the first two principal components is shown in parentheses; symbols indicate sampling locations of populations and colors correspond to the six genetic clusters assigned to each individual by Bayesian clustering for isolates with ≥0.5 membership to a single cluster; grey symbols are isolates not assigned to any cluster when using the threshold of 0.5. (**C**) Delta K plot of the STRUCTURE Bayesian clustering analyses for Ks of 2–12. (d) Bar plots of membership proportions (y-axis) as inferred by STRUCTURE for K=2, K=6 and K=7; each isolate displayed as a vertical bar; each color represents a cluster and isolates are ordered according to longitude (West to East); sample sizes are: MdR=33, Bret=4, Kew=14, Auv=27, Cev=31, Aub=22, AubX=16, El=26, So=22, Other1-4=3+3+6+4. For code and origin of populations see Table 1.

**FIGURE 3.**
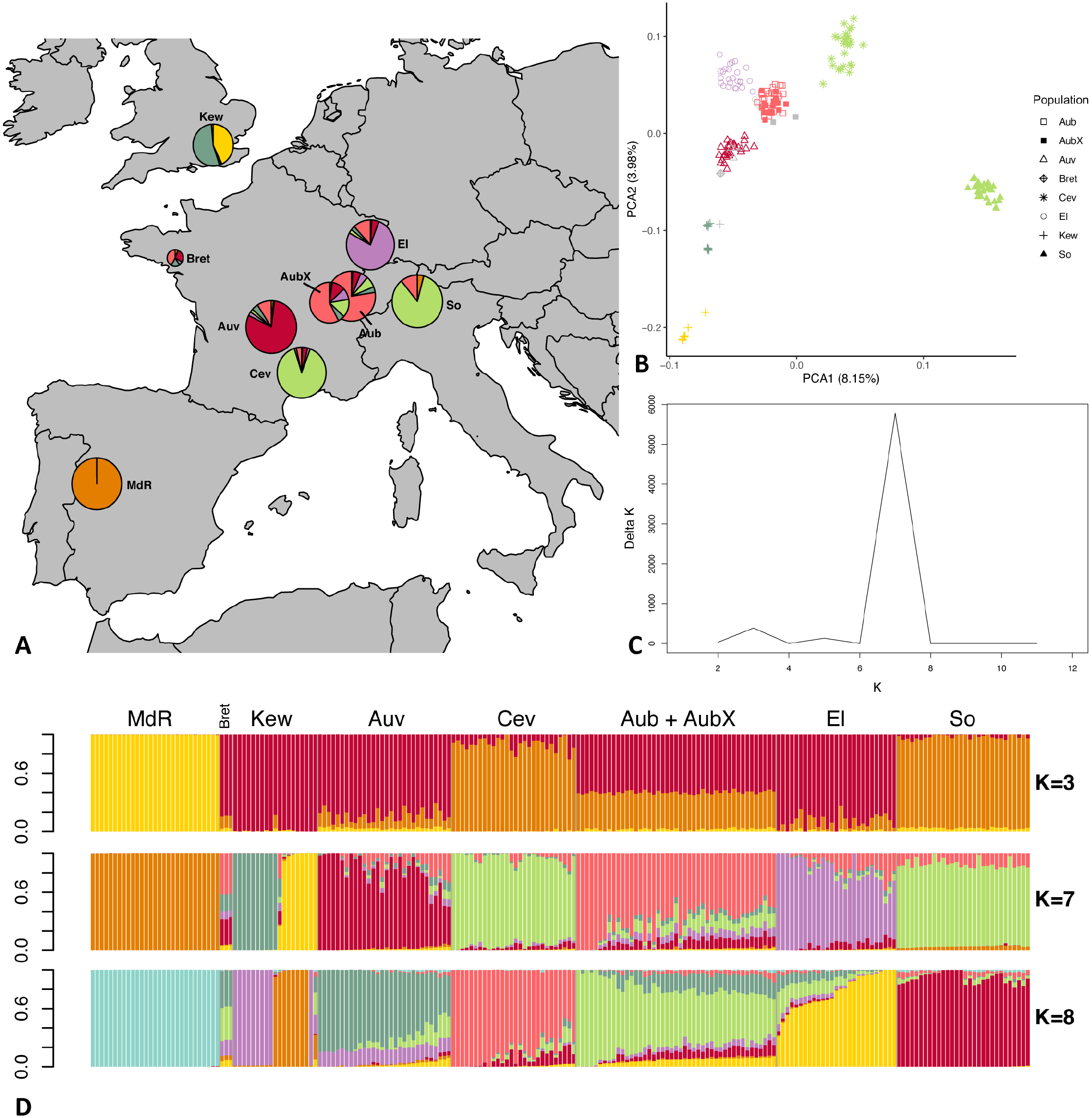
Population structure analyses of 211 *E. clarkii* isolates based on genomewide SNPs. The LD pruned dataset including 79,352 SNPs was used in the STRUCTURE analyses whereas the complete dataset of 400,033 SNPs was used for the PCA. (**A**) Sampling locations of *E. clarkii* individuals infecting *H. lanatus* and *H. mollis* (Kew & Bret); pie charts represent Bayesian cluster membership proportion for K=7, diameters reflect sample size (range n=3 to n=30). (**B**) Principal component analysis of genetic differentiation among isolates excluding the population from Spain (MdR); The percentage of variance explained by the first two principal components is shown in parentheses; symbols indicate sampling locations of populations and colors correspond to the seven genetic clusters assigned to each isolate by Bayesian clustering for isolates with ≥0.5 membership to a single cluster; grey symbols are isolates not assigned to any cluster when using the threshold of 0.5. (**C**) Delta K plot of the STRUCTURE Bayesian clustering analyses for Ks of 2–12. (d) Bar plots of membership proportions (y-axis) as inferred by STRUCTURE for K=3, K=7 and K=8; each isolate displayed as a vertical bar; each color represents a cluster and isolates are ordered according to longitude (West to East); sample sizes are: MdR=29, Bret=3, Kew=19, Auv=30, Cev=28, Aub=27, AubX=18, El=27, So=30. For code and origin of populations see Table 1.

Consistent with results obtained with PCA, STRUCTURE showed evidence of subdivision according to geography in both species. The Spanish population was assigned to a distinct cluster (K) irrespective of the inferred number of clusters (K=2 to K=12), and there was no genetic differentiation between isolates sampled at different time points in the population from Aubonne (AubX and Aub sampled 2005 and 2018). In *E. typhina* the most likely number of clusters was K=6 as inferred by the maximized likelihood and ΔK values across twelve replicate runs (Figure 2A,C,D). At K=6, two of the six clusters contained all isolates from a single location (red: MdR, green: So). Isolates from most locations appeared to have ancestry from the same orange cluster, to varying degrees. Isolates from more Northern locations (El, Kew, Bret) and three isolates from Plymouth showed the highest ancestry proportion to this cluster. The exception was isolates from Cev, the majority of which had genotypes assigned to two distinct clusters (yellow and darkgreen) but also contained admixed isolates. Additionally, the population from Western Switzerland contained seven isolates assigned to a sixth cluster (purple), three of which were sampled in 2005 (AubX) and four in 2018 (Aub) (see Supplementary Figure S7), but the majority of isolates had mixed ancestry. A number of isolates from Aub and Cev as well as six additional isolates collected in the Southeastern tip of France showed a higher proportion of mixed ancestry with membership proportion <0.5 for any of the inferred clusters (grey symbols in PCA). The relationship among genetic clusters and location did not get much clearer at K=7.

In *E. clarkii* the most likely number of clusters inferred by STRUCTURE was K=7 (Fig 3A,C,D). At K=7 all clusters consisted of isolates from a single sampling location except for the populations from Southern France (Cev) and Southern Switzerland (So), which were assigned to the same cluster. At lower and higher Ks (K=5, K=6, K=8, Supplementary Figure S8), however, these two populations were assigned to distinct clusters. The population collected at Kew Gardens (Kew) consisted of genotypes from two distinct clusters as well as a single admixed isolate. Although Central European populations contained a higher number of individuals with mixed ancestry, admixture proportions were lower than in *E. typhina*. Overall, ten isolates were not assigned to a cluster as they showed <0.5 membership proportion to any single cluster. Of these admixed isolates, all except for three genotypes collected in Western France (Bret) and the admixed isolate from Kew shared the majority of their ancestry with their population of origin.

Results from DAPCs were consistent with STRUCTURE analyses suggesting seven clusters (corresponding to geographic locations) a best-fit in *E. clarkii*. In *E. typhina* DAPC suggested an optimal K of 3 with MdR (Spain) as the most distinct cluster and the other populations separating on the second axis. When removing MdR, K=2 was optimal dividing genotypes into a Northern and a Southern cluster. At higher K’s the remaining six populations could be distinguished with corresponding to geographic locations and with admixture among the northern and central populations.

### 3.2 Genetic differentiation between populations and isolation by distance

Weighted genome-wide averages of pairwise FST across all loci were lower between *E. typhina* populations (0.05-0.21, Table 2), supporting higher levels of gene flow, particularly among Central European populations that the clustering analysis inferred mixed ancestry (Kew, Auv, Aub and El). *E. clarkii* populations had higher values of F_ST_ consistent with lower levels of gene flow (0.12-0.49, Table 2). Population pairs with the highest F_ST_ values also had the most fixed differences between them, and overall, there were more fixed differences between *E. clarkii* populations than *E. typhina* (Supplementary Table S4). We found a positive correlation between genetic differentiation and geographic distance consistent with a pattern of isolation-by-distance (IBD) in both species (Figure 4). A comparison of the slopes of the two linear models showed that the effect of IBD was stronger in *E. clarkii* compared to *E. typhina* with higher genetic differentiation between population pairs [Z= 6.964, p= 3.314e-12, ‘lm_slopes_compare’ from R package EMAtools (Kleiman 2017)].

**FIGURE 4.**
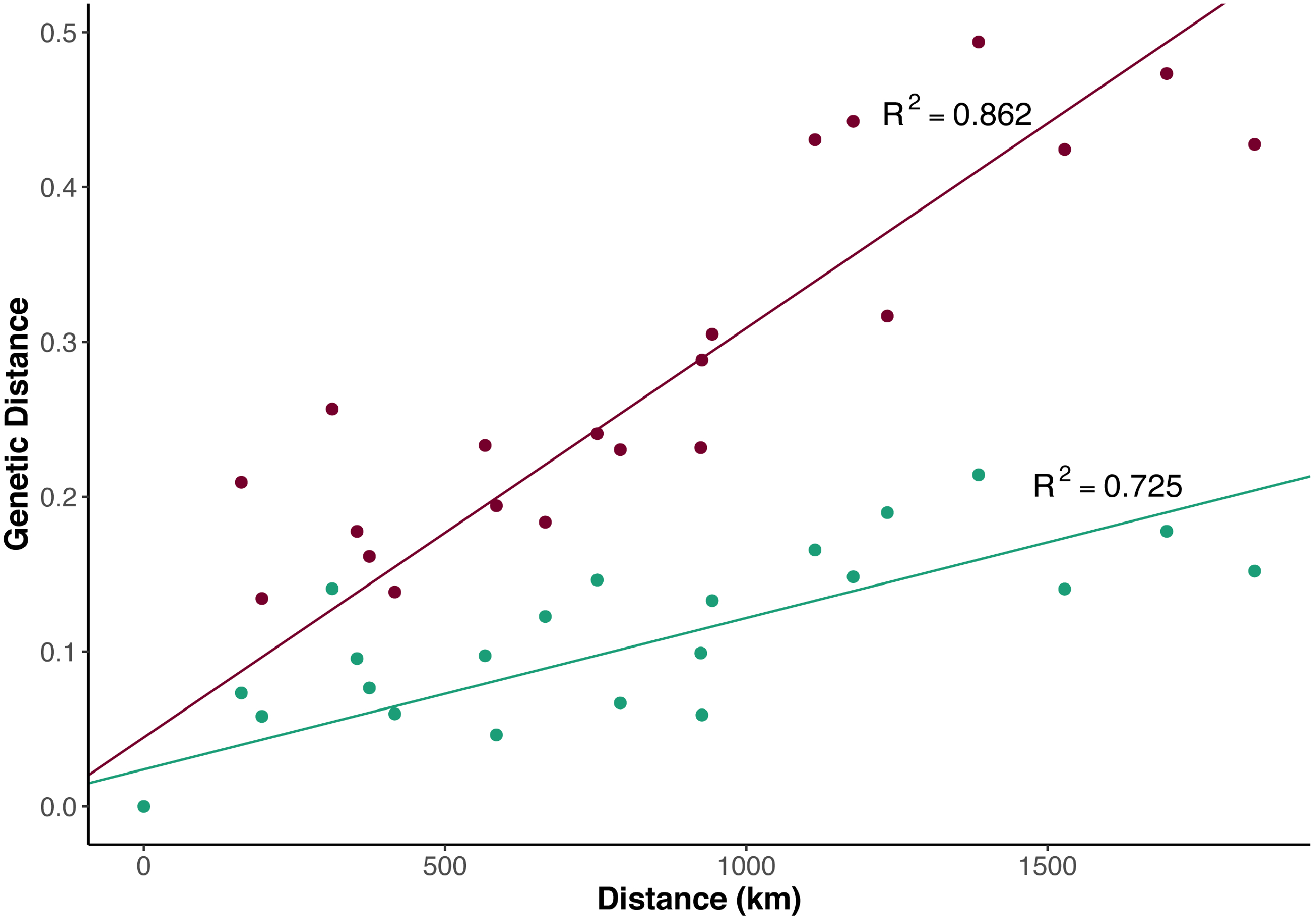
Isolation by distance among *Epichloe* populations. Relationship between pairwise *F*_ST_ and geographic distance for *E. typhina* (green) and *E. clarkii* (purple). Correlations for both species were significant (*E. typhina:* p= 4.19e-08; *E. clarkii:* p= 7.187e-13) with increasing geographic distance having a stronger effect in *E. clarkii* (p= 3.314e-12).

**TABLE 2.**
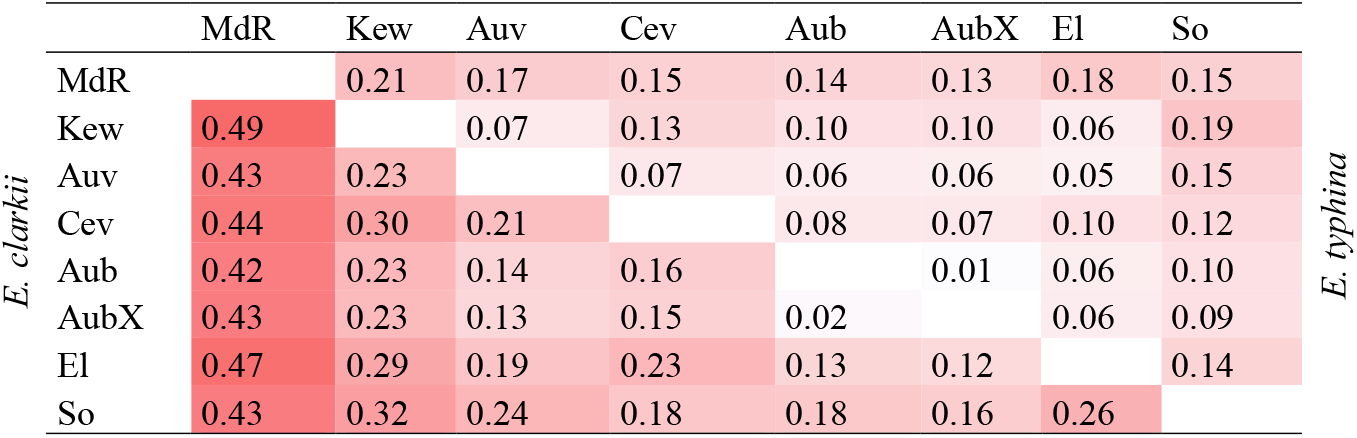
Mean genome-wide *F*_ST_. Pairwise weighted *F*_ST_ values (Weir and Cockerham, 1984) for populations of *E. typhina* (top right) and *E. clarkii* (bottom left), calculated using genome-wide SNPs. Populations are ordered according to longitude (West to East) and higher *F*_ST_ values are underlaid with darker shades of red.

### 3.3 Genetic diversity within populations

Mean genome wide nucleotide diversity (π) ranged from 0.0024 to 0.0027 in *E. typhina* populations and from 0.0008 to 0.0012 in *E. clarkii* populations (Supplementary Figure S9, Supplementary Table S5). This was consistent with a previous study which found higher diversity in *E. typhina* compared to *E. clarkii* in one sympatric population pair (Schirrmann *et al*. 2018).

Mean Tajima’s D was slightly negative in all populations of *E. typhina*, ranging from −0.369 to −0.061 (Supplementary Figure S10, Supplementary Table S6). In *E. clarkii* Tajima’s D showed more variation across populations: three populations, including the same population sampled at different time points (Aub and AubX), had slightly negative Tajima’s D ranging from −0.354 to −0.015, two populations had slightly positive Tajimas’s D (Auv = 0.102 and Cev = 0.151) and two populations had Tajimas’s D around 1 (Kew = 0.903 and MdR = 1.318).

### 3.4 Analysis of linkage disequilibrium

Linkage Disequilibrium (LD) decayed quickly with distance, r^2^ <0.2 in 1-2kb in all *E. typhina* populations and most *E. clarkii* populations (Supplementary Figure S11). Two populations in *E. clarkii* were exceptions/distinct (Supplementary Figure S12): In Kew, LD decayed more slowly with distance, which is most likely a result of clonal population substructure. The Spanish population MdR had very high levels of LD overall, which could indicate a recent bottleneck.

### 3.5 Effective population size *N_e_*

We used Watterson’s estimators of θ (Watterson 1975) and an approximate mutation rate of 3.3 × 10^-8^ per site to calculate and compare effective population sizes within and between *Epichloe* species. Effective population sizes in *E. typhina* were between 2-fold and 4-fold larger than in *E. clarkii* populations, which also had more variance in Ne estimates among populations (Table 3: Estimates of effective population size *N*_e_). In *E. typhina* the population from Western Switzerland (Aub) had the largest *N_e_* (43,941) and the British population (Kew) had the smallest (37,034). In *E.clarkii* the population from Southern Switzerland (So) had the largest *N_e_* (18,809) and the Spanish population (MdR) had the smallest (9,251).

**TABLE 3.** Effective population size *N_e_* of different populations of *E. typhina* and *E. clarkii*.

| Population<sup>1</sup> | *E. typhina* | *E. clarkii* |
| --- | --- | --- |
| Aub | 43,941 | 17,606 |
| AubX | 41,997 | 18,010 |
| Auv | 42,358 | 15,284 |
| Cev | 40,223 | 15,989 |
| El | 41,235 | 14,289 |
| Kew | 37,034 | 10,250 |
| MdR | 39,535 | 9,251 |
| So | 39,647 | 18,809 |

<sup>1</sup>For code and origin of populations see Table 1.

## 4 Discussion

Our parallel study of two closely related and ecologically similar fungal pathogens reveals how sympatric species can experience distinct evolutionary contexts across a shared geographical range. Despite the two species sharing a highly specialized host-sterilizing lifestyle, we identified striking differences in their genetic differentiation, genetic variation and population size. In the following these findings are discussed in the context of the phylogeographic history, pathogen biology, and selection mediated by interactions with the host.

### 4.1 Different levels of gene flow among spatially structured populations

In our analysis of genome-wide variation we found a clear pattern of biogeographic structure in both *E. typhina* and *E. clarkii*. Spatially separated populations were also genetically differentiated, and we were able to clearly resolve both the West-to-East and South-to-North distribution of our sampling. In both species genetic distance increased with geographic distance and the most remote population from Spain was genetically very distinct. Despite the similar genetic structure in *E. typhina* and *E. clarkii* at the European scale, patterns of geneflow among populations are different. *E. typhina* populations had higher genetic diversity and showed less differentiation between populations indicating greater gene flow particularly among Central European and Northwestern populations. In contrast, we found lower diversity and higher levels of genetic differentiation among *E. clarkii* populations and a stronger effect of isolation by distance. Comparison of genome organization and size between the reference genomes of the two species suggested that genome expansion in *E. clarkii* compared to *E. typhina* may be linked to a reduction in effective population size leading to stronger effects of drift compared to *E. typhina* (Treindl *et al*. 2021). The lower genetic diversity and lower effective population size within *E. clarkii* populations we report here supports this hypothesis.

Of the total variation (number of SNPs) assessed among populations in both species, a similar proportion was polymorphic within most populations indicating that the two species likely have similar mutation rates and that, in spite of a lower overall diversity, *E. clarkii* populations still harbor a considerable amount of standing genetic variation. This can be expected given the obligately outcrossing lifestyle of the pathogen and high recombination rates detected. The exceptions were two populations, Kew and MdR, which were found to have significantly less genetic diversity and high degrees of linkage likely related to population substructure and/or a recent bottleneck which is also reflected in the positive Tajimas’D values for these two populations. Such demographic processes affecting genomewide variation will need to be taken into consideration in future studies when investigating the impact of selection in these populations.

### 4.2 Determinants of population structure in *Epichloe* pathogens

#### 4.2.1 Phylogeographic history may shape genetic structure

Genetic variation in specialized obligate pathogens such as *E. typhina* and *E. clarkii* is expected to be strongly influenced by the distribution and genetic structure of their host plants (Wilson *et al*. 2005; Greischar and Koskella 2007). Previous studies of genetic structure in the anther smut pathogen *Microbotryium lynchnidis-dioicae* and its white campion host (*Silene latifolia*) across a larger sampling range have identified three genetically differentiated clusters spanning greater geographic scales and suggested that clusters reflect the concerted recolonization of the host plant and its fungal pathogen from distinct southern glacial refugia (Vercken *et al*. 2010; Feurtey *et al*. 2016; Badouin *et al*. 2017). The maintenance of genetically differentiated gene pools in these refugia and subsequent northwards recolonization as ice-sheets retreated is thought to have strongly impacted the spatial structure of genetic variation observed across Europe today and indeed, the existence of at least three major ice-free refugia during the last glacial maximum located in Iberia, Italy and the Balkans is a well-established hypothesis supported by phylogeographic studies of a growing number of temperate species including grasses (Hewitt 2000; Provan and Bennett 2008; François *et al*. 2008). For example, recent studies found genetic structure of natural populations to be consistent with postglacial colonization history in the *Poaceae* grasses *Festuca pratensis* (Fjellheim *et al*. 2006), *Festuca rubra* (von Cräutlein *et al*. 2019) and *Lolium perenne* (McGrath *et al*. 2007). Genetic diversity in Europe has also been assessed in one of our host grasses, *D. glomerata*, and three genetic clusters could be identified although genetic differentiation was low among natural populations at large geographic scales across Europe (Tuna *et al*. 2004; Last *et al*. 2013). The relatively high levels of genetic diversity and genetic admixture reported for the species in these studies are not surprising given that for outcrossing and wind pollinated grass species gene flow among populations is expected to be high (Brown 1978). Furthermore, human mediated transfer likely also plays a role in promoting genetic admixture in grasses which are also used as cultivars in farming (Last *et al*. 2013; von Cräutlein *et al*. 2019). Although the phylogeographic origin has likely influenced genetic structure of host populations and this may be reflected in the genetic structure of *E. typhina* and *E. clarkii*, analysis of a larger geographic area ideally spanning the entire distribution range and including Eastern and Southern European populations would be necessary to be able to detect patterns of glacial refugia. The data in this study represents a smaller scale of sampling and the fine-scale population structure of the two pathogens suggests that additional processes play a role in driving genetic differentiation between populations.

#### 4.2.2 Distribution and dispersal of pathogens and hosts

Both *D. glomerata* and *H. lanatus* are native and widely distributed plants in grasslands across Europe and share similar long-distance pollen dispersal mediated by wind whereas seeds usually disperse more locally (Düll and Kutzelnigg 2016). Our sampling locations are situated in natural grassland ecosystems where the impact of anthropogenic use is likely minimal. Nevertheless, it is possible that agricultural practices such as sowing for pasture farming or forage production could have differentially affected the genetic make-up of grass populations in the two host species. While *H. lanatus* is not commercially sown due to its low nutritional value as a forage grass and may even be considered a weed, *D. glomerata* cultivars are frequently sown as part of commercial forage grass mixtures. Outcrossing of such cultivar genotypes into natural populations may promote gene flow and have a homogenizing effect on genetic structure in *D. glomerata*, however, we expect admixture to be reduced by the fact that pastures are usually mown before grasses produce flowers or seeds. We have not assessed the underlying genetic structure of natural host populations in this study, however, higher levels of gene flow and reduced genetic differentiation among *D. glomerata* populations could explain why its pathogen *E. typhina* also shows more admixture and higher diversity. Populations of *H. lanatus* on the other hand are unlikely to experience more gene flow due to anthropogenic practices and populations may therefore remain more differentiated which is reflected in the genetic structure of *E. clarkii*. While both host grass species are similarly widespread across our study area and often co-occur in the same habitat, the abundance of their obligate pathogens differs: *E. clarkii* can be very common locally, such is the case of sympatric populations sampled for this study, but generally has a patchier distribution whereas infections of *E. typhina* on *D. glomerata* are more widely distributed. This pattern is consistent with the genetic data, supporting the smaller population sizes and lower levels of gene flow among *E. clarkii* populations. We hypothesize that the different spore sizes of *E. typhina* and *E. clarkii* differentially impact the dispersal abilities of these pathogens. *E. typhina* produces long, thread-like spores which may entail more efficient long-distance dispersal by wind whereas *E. clarkii* produces shorter and wider part-spores (White Jr. 1993; Leuchtmann and Clay 1997). The resulting large number of part-spores may be more efficiently dispersed over shorter distances and increase disease pressure locally.

#### 4.2.3 The role of local adaptation

Despite the abundance of host plants and the large production of wind-dispersed meiotic spores in both *E. typhina* and *E. clarkii* which enable contagious spread to new host plants (Leuchtmann and Clay 1997)*, Epichloe* infections are not everywhere. The strong pattern of genetic structure suggests that effective long-distance dispersal may be limited, for example by the range of susceptible host genotypes. For obligate biotrophic fungi such as *Epichloe* that depend on their living host, the host plant constitutes the distinct ecological niche that the pathogen evolves in. The ongoing coevolutionary arms-race in such host-pathogen systems is thought to be one of the most important drivers of natural selection acting at the population level and is generally assumed to result in local adaptation of pathogen populations to host genotypes and vice-versa (Greischar and Koskella 2007). This may be reflected in congruence of genetic structure between host and pathogens (Feurtey *et al*. 2016; Hartmann *et al*. 2020), however, the extent to which co-evolutionary processes lead to genetic co-structure and local adaptation strongly depends on the strength of selection exerted by both partners, dispersal abilities related to levels of gene-flow and the role of local environmental factors (Croll and McDonald 2016). As discussed, levels of gene flow for outcrossing and wind pollinated grass species such as *D. glomerata* and *H. lanatus* are expected to be high, nevertheless, the abundance of rare alleles specific to geographically distinct populations supports the assumption that *D. glomerata* populations may be locally adapted (Tuna *et al*. 2004; Last *et al*. 2013). For *H. lanatus* genetic diversity has, to our knowledge, not been assessed to date, however, reciprocal transplant experiments provided some evidence for local adaptation of European *H. lanatus* populations (Bischoff *et al*. 2006). Given that *E. typhina* and *E. clarkii* are highly specialized pathogens that cannot grow or reproduce without establishing a systemic infection in their host plant, it is expected that co-evolutionary processes have shaped the fine-scale genetic structure in this system and that local adaptation may have contributed to the accumulation of genetic differences among *Epichloe* populations. Future studies should aim to further investigate patterns of local adaptation by assessing congruence of genetic structure in these two host-pathogen pairs combined with empirical tests: If pathogen populations are locally adapted they should be more fit on their local host genotypes than on foreign host genotypes (home vs away) and should outperform foreign pathogen genotypes on local hosts (local vs foreign) (Croll and McDonald 2016). This could be assessed experimentally in the *Epichloe* study system by performing reciprocal inoculation experiments with plants from natural populations.

### 4.3 Additional host species confirmed for *E. clarkii*

In our extensive sampling of natural populations of *E. clarkii*, we found two locations in the UK and in Bretagne, Northern France where the pathogen occurred on the host grass *Holcus mollis*. This was surprising given that *E. clarkii* is considered to be highly specialized on *H. lanatus* and, moreover, *H. mollis* is the host of a distinct *Epichloe* species, *E. mollis* (Leuchtmann et al. 2014). However, a literature review revealed that this finding indeed backs up previous observations from the UK, where *E. clarkii* was identified on *H. mollis* based on its ascospore morphology (Spooner and Kemp 2005). We now provide additional evidence with genetic data and, for the first time, report *H. mollis* as an *E. clarkii* host on European mainland. It needs to be noted that hybrids between *H. lanatus* and *H. mollis* have been described (*Holcus x hybridus* Wein, which usually show morphologically intermediate phenotypes but tend to resemble *H. mollis* more closely (Beddows 1961; Carroll and Jones 1962). In our samples, species identification of the putative *H. mollis* host grasses was based on the presence of orthotropic shoots/rhizomes indicating clonal propagation and the presence of longer hairs on the nodes compared to *H. lanatus*. Characters of the inflorescence could not be considered since they were encased in the fungal fruiting structure. Although these morphological characters match well with descriptions of *H. mollis*, we cannot entirely exclude that samples originate from hybrids, and this will need to be investigated further. In any case this finding opens up new avenues to investigate the evolution of host specialization and divergence within the *Epichloe* study system.

## 5 Conclusion

The comparison of genetic structure within and between populations of *E. typhina* and *E. clarkii* indicate distinct evolutionary histories of the two species and this is likely influenced by biological factors such as dispersal ability as well as the underlying genetic structure of host populations which may differentially affect coevolutionary dynamics between the two pathogens and their hosts. It appears that *E. typhina* infecting *D. glomerata* is a more ubiquitous pathogen with higher dispersal abilities among more genetically homogenous host populations and *E. clarkii* infecting *H. lanatus* and *H. mollis* forms more fragmented populations restricted by lower dispersal abilities and adaptation to local host genotypes. We expect that in both systems, sufficiently low levels of gene flow and divergent selection should promote local adaptation to the host, however, *E. clarkii* with its lower effective population size is more likely to experience metapopulation dynamics with frequent local extinction and recolonization events which may lead to loss of genetic diversity within populations and generate genetic differences between populations through bottlenecks and genetic drift. Future work should investigate signatures of adaptive variation in these two pathogens in order to disentangle to effects of drift and selection. This study and dataset presented here represents an essential basis for understanding coevolution and adaptation in the *Epichloe* study system and providing novel insights into the population structure and patterns of gene-flow. It demonstrates how the biology of these species and the interaction with their host plants may influence evolutionary trajectories and helps to increase our understanding of the population genetic processes that generate and maintain diversity of fungal pathogens in natural ecosystems.

## Author contributions

AL and AT conceived the study design. AT and JS developed methodology and did the formal analyses. AT wrote the initial draft. AT, JS and AL reviewed and edited the manuscript. AL acquired funding. All authors have read and agreed to the submitted version of the manuscript.

## Acknowledgements

We would like to kindly thank Martyn Ainsworth (Royal Botanical Gardens Kew, UK), Jean-Paul Priou, James M. Neenan, Elisabeth Walder and Lilith Treindl for assistance with sample collection, and Claudia Michel and Beatrice Arnold for laboratory assistance. Fabrizio Steinebrunner provided samples for DNA extraction from a previous collection at the Aubonne site. We thank Karsten Rohweder for help with the GPS device and Niklaus Zemp for bioinformatic support. Data presented and analyzed here were generated in collaboration with the Genetic Diversity Centre (GDC), ETH Zurich and the Functional Genomics Center Zurich (FGCZ), UZH.

## Data availability statement

Data associated with this manuscript will be deposited in the Sequence Read Archive and made publicly available upon publication.

## Supplementary material

The Supplementary material for this article can be found online at the publisher’s website:

Supplementary Notes

Table S1 SNP datasets used for different analyses

Table S2 Number of segregating sites per population

Table S3 Proportion of segregating sites per population [%]

Table S4 Fixed differences between populations

Table S5 Mean nucleotide diversity (π) per site

Table S6 Mean Tajimas D calculated over 40kb windows

Figure S1 Defining filtering cutoffs

Figure S2 Summary of statistics after filtering

Figure S3 PCAs based on SNP datasets with different filtering criteria

Figure S4 PCAs based on SNP datasets with no missing data

Figure S5 PCAs within populations of *E. typhina*

Figure S6 PCA within populations of *E. clarkii*

Figure S7 *E. typhina* STRUCTURE results for all Ks

Figure S8 *E. clarkii* STRUCTURE results for all Ks

Figure S9 Nucleotide diversity per population

Figure S10 Tajima’s D per population

Figure S11 Decay of linkage disequilibrium (r^2^) with physical distance in *E. typhina* and *E. clarkii* populations

Figure S12 Decay of linkage disequilibrium (r^2^) with physical distance in *E. clarkii* Kew and MdR

Figure S13 PCAs by genome compartments in *E. typhina*

Figure S14 PCAs by genome compartments in *E. clarkii*

Figures S15 and S16 Population summary statistics by genome compartments

Figure S17 Proportion of mating types in *E. typhina* populations and across all sequenced genotypes

Figure S18 Proportion of mating types in *E. clarkii* populations and across all sequenced genotypes

