## Supplementary Notes for "Genetic diversity and population structure of *Epichloe* fungal pathogens of plants in natural ecosystems"

#### Variant filtration

The use of Rank Sum Tests implemented in GATK (McKenna et al. 2010), is commonly applied in many variant filtration pipelines, including haploid datasets (e.g. Hartmann et al. 2018, Mohd-Assaad et al. 2018). However, RankSum annotations using the GATK VariantFiltration tool are only calculated for heterozygous sites with read support for both the REF and ALT allele. In haploid individuals true variants should only contain reads for either the REF or ALT allele and erroneously "heterozygous" sites with support for both alleles are evidence of sequencing errors or misalignments during mapping. While one of the called alleles may be real, the others are a result of mapping artifacts or sequencing errors and therefore RankSumTest statistics should be positively biased towards the "real" allele. Based on these considerations we have not used and do not recommend using Rank Sum Test statistics (calculated within individuals) for filtering of haploid datasets. For further information see details on ReadPosRankSumTest, MQRankSumTest and BaseQRankSumTest here: https://gatk.broadinstitute.org/hc/en-us/categories/360002369672-Tool-Index and notes on hard filtering here: https://gatk.broadinstitute.org/hc/en-us/articles/360035531112--How-to-Filter-variants-either-with-VQSR-or-by-hard-filtering.

The filtering step to exclude SNPs with >30% missing data within one or more populations was performed using only the eight sympatric populations and not for the additional allopatric samples which only included few isolates per locality.

#### Datasets

After filtering we retained 658,021 and 400,033 high-quality biallelic SNPs in the *E. typhina* and the *E. clarkii* dataset respectively. For downstream analyses we applied additional filters to subset this dataset: To calculate popualtion statsitics we removed isolates that were sampled from allopatric populations and removed variants from the SNP dataset that were monomorphic among populations. We also generated a dataset containing only SNPs genotyped across all isolates (no missing data). For the STRUCTURE analysis, we applied a minor allele frequency filter of 0.05 and LD-pruned variants for sites with a pairwise genotypic r2 greater than 0.3 in 10kb windows. For DAPC, this dataset was additionally thinned to one variant per 2kb.

**Replicates**

We included technical replicates of five and three individuals respectively. For each replicate, we removed the replicate with fewer mapped reads from the dataset after assessing genotype concordance rates. Average allele concordance between replicates was 98.9% with a standard deviation of 0.1% in Et and 97.7% with a standard deviation of 0.3% in Ec.

**Population sub-structure**

We investigated genetic sub-structure within sympatric populations of *E. typhina* and *E. clarkii* to identify highly related individuals using PCA as described in the main text. Most populations did not have apparent sub-structure or contain highly related individuals. The exceptions in *E. typhina* were the populations Cev, with 5 individuals that were more distinct from the rest in principal component space, and MdR with 2 clearly separated genotypes. In *E. clarkii* the Kew population showed a high degree of sub-structure with three groups of highly related individuals and a single intermediate genotype. Individuals from each sub-populations corresponded to different areas within the sampling location and the very low level of diversity within sub-populations compared to the whole population suggested that these may indeed be isolates of clonal origin. This was not highly surprising and likely related to the clonal population structure of the host grass *H. mollis.*

**PCA analysis and calculation of summary statistics by compartments**

*Epichloe* genomes are compartmentalized into conserved gene-rich regions and AT-rich regions containing a high proportion of repeat elements and only few genes (Chapter I of this thesis). Accurate variant discovery in these polymorphic AT-rich regions using alignment of short-reads is often challenging because reads are more likely to map incorrectly (see Pfeifer 2017). We used stringent filtering criteria to remove erroneous SNP calls and assessed the distribution of variants in our filtered dataset. For this, genome-wide SNPs (complete datasets, see Supplementary Table S1) were split by compartment to generate two variant files for each species: one containing SNPs called in gene-rich regions and one containing the SNPs called in AT-rich regions. The number of SNPs within each compartment was roughly proportional to the relative size of the compartment: In *E. typhina* the AT-rich compartment made up 31.7% of the genome and contained 23.9% of SNPs whereas in *E. clarkii* the AT-rich compartment made up 48.6% and contained 53.1% of SNPs. We used these SNP datasets to analyze genetic structure among populations using PCA and found that clusters corresponding to sampled populations were generally resolved with variants in either compartments giving us confidence that our SNP datasets reflect true biological variation (Supplementary Figures S13 and S14). However, genetic differentiation between clusters was more apparent in the gene-rich compartment and more variance was explained by these variants in both species. In *E. typhina* clusters were less clearly resolved in the AT-rich compartment which may suggest lower genetic differentiation in these regions of the genome (Supplementary Figure S13 C & E). The overall pattern of population structure was more strongly shaped by strong genetic differentiation in the gene-rich regions and the high number of SNPs there. In *E. clarkii* genetic differentiation between Cev and So was less pronounced in the AT-rich compartment compared to the gene-rich, which could explain why these populations were assigned to the same cluster in the overall STRUCTURE analysis at K=7. We hypothesize that the patterns of genetic differentiation in distinct genome compartments could reflect the differential action of random genetic drift and selection in these genome compartments: Variation in the conserved gene-rich regions may be strongly influenced by local selection driving differentiation between populations while the AT-rich compartment evolves under a stronger influence of drift.

We also computed the summary statistics $\pi$, Tajima's D, LD $r^2$ and SNP density for each compartment separately using the same parameters as in the genome-wide analysis. Compiled plots for each population can be found in the Supplementary (Figures S15 and S16). Across all populations, rare alleles were more abundant in the AT-rich compartment (lower Tajima's D than in the gene-rich compartment) and this could reflect the activity of RIP (repeat-induced point mutation) in these regions of the genome (most mutations introduced by RIP are expected to be at low frequency unless they are targeted by selection). Furthermore, SNPs were more strongly linked in the gene-rich compartment. This difference was most striking in *E. clarkii* MdR where the gene-rich compartment had an extremely high mean $r^2$ consistent with the idea that this population may have experienced a recent bottleneck. Average SNP density in 10kb windows was generally lower in AT-rich regions and this is likely influenced by the fact that many variants in regions were mapping was very poor were removed during filtering resulting in a relatively large number of "empty" AT-windows that did not contain any more SNPs. Nevertheless, outlier points indicate that we also retained a number of highly polymorphic windows with higher-than-average SNP density.

**Calculation of effective population size**

To compare effective population sizes within and between species we estimated effective population sizes ($N_e$) from polymorphism data with the parameter $\theta = 2N_e\mu$. Calculations were based on Watterson's estimators of θ (Watterson 1975), which is based on the number of segregating sites in a population relative to the sequence length, and an assumed mutation rate (μ) of $3.3 \times 10^{-8}$ per site. For each population we calculated the number of segregating sites in 5kb windows along the genome using the PopGenome R package (Pfeifer et al. 2014). We divided the number of segregating sites by the $(n-1)^{th}$ harmonic number to calculate Watterson's θ, with n being the number of chromosomes or in case of haploids simply the sample size. Following the formula, we then calculated $N_e = \theta/2\mu$.

**Assessment of hybridization in sympatric populations**

A previous study by Bultman et al. (2011) found that 9.3 % of fruiting bodies collected from a sympatric population of *E. typhina* and *E. clarkii* contained asci with hybrid spores. We assessed the occurrence of hybridizations in two additional natural populations in Southern and Central France (Cev and Auv) one year after the collection of population samples. Populations were revisited at the end of the flowering season in July 2018 and 1-3 ripe stromata with perithecia of *E. typhina* and *E. clarkii* were collected from 7-13 *D. glomerata* and *H. lanatus* plants. From each stroma three probes were taken and ascospore morphology within perithecia was observed microscopically. When asci contain the distinct spore-morphotypes of both *E.typhina* and *E.clarkii,* this indicates that hybridizations have occurred (see Treindl and Leuchtmann 2019). We detected hybrid asci in 12.5 and 8.3% (mean 10.4%) of probes taken from naturally fertilized fruiting bodies (stromata) collected from sympatric populations in Southern and Central France (Cev and Auv, Supplementary Table S9-S11). Despite the occurrence of interspecific mating, however, we did not detect any intermediate/ putatively hybrid genotypes in a preliminary analysis using a draft version of the *E. typhina* reference genome and individuals from both species, suggesting lack of gene flow between *E. typhina and E. clarkii*. Additionally, we did not find evidence for hybrid or introgressed isolates infecting either *D. glomerata* or *H. lanatus* despite the extensive analyses of over 400 isolates from sympatric populations (where the potential for hybridization exists). These observations suggest that despite incidences of interspecific mating as evidence by the occurrence of hybrid asci in sympatric populations, *E. typhina* and *E. clarkii* are reproductively isolated and that postzygotic reproductive barriers exist that prevent the establishment of hybrid genotypes in the population and maintain the genetic and ecological integrity of the two species.

### Mating types

We analyzed the distribution of mating types in sympatric populations as skewed sex ratio within populations reduces effective population size ($N_e$) and can bias population genomic inferences (Wright 1938). Therefore, we analyzed the distribution of mating types in sympatric populations. Sexual *Epichloe* species have a bipolar heterothallic mating system (White and Bultman 1987). Each haploid individual possesses a mating type locus with one of two alternative idiomorphs designated MTA and MTB or mat-1 and mat-2. The MTA idiomorph includes three genes (mtAA, mtAB, and mtAC) whereas the MTB idiomorph includes only the gene mtBA (Schardl et al. 2012). The reference genomes for both species have the MTA mating type idiomorph (mat-1) so to assess sex ratios in our populations (i.e. the proportions of mating types) we extracted the region including the mating type locus from the alignment files (bam) of every individual and used the function genomecov -bga in *bedtools* version 2.28.0, to report read depth in a BedGraph format (Quinlan and Hall 2010). This includes regions with zero coverage and can be visualized in IGV (Integrative Genomics Viewer) (Robinson et al. 2011). Individuals with the same mating type as the reference (mat-1) have normal read coverage whereas individuals with the alternative mating type (mat-2) have a deletion/ no read coverage. Here we included all individuals with sufficient coverage around the locus, even those with low coverage overall that were removed for subsequent analyses. We assessed the proportions of mating types and found that both mating types were present in all sampled populations. Across all individuals in both *E. typhina* and *E. clarkii* sex ratios were slightly biased towards mating type 1 with 54% in *E. typhina* and 55% in *E. clarkii* (Supplementary Figures S17 and S18). Among populations sex ratios varied from 41% to 68% mat-1 in *E. typhina* and 39% to 71% in *E. clarkii*. This variation is likely to be increased by stochastic processes due to small sample sizes.
