## Supplementary Tables & Figures for "Genetic diversity and population structure of *Epichloe* fungal pathogens of plants in natural ecosystems"

**Supplementary Table S1.** SNP datasets used for different analyses.

| | *E. typhina* SNPs | *E. clarkii* SNPs | Analyses |
|---|---|---|---|
| complete | 658,021 | 400,033 | PCA |
| populations only | 657,966 | 397,283 | population statistics |
| no missing data | 295,446 | 63,930 | PCA |
| LD-pruned | 152,883 | 79,352 | PCA, STRUCTURE |
| LD-pruned and thinned | 12,562 | 11,338 | DAPC |
| complete gene-rich | 500,747 | 187,711 | PCA |
| complete AT-rich | 157,274 | 212,322 | PCA |

**Supplementary Table S2.** Number of segregating sites per population.

| | *E. typhina* | *E. clarkii* |
|---|---|---|
| Aub | 362,128 | 206,283 |
| AubX | 317,031 | 189,525 |
| Auv | 368,058 | 183,846 |
| Cev | 361,706 | 189,053 |
| El | 354,895 | 167,418 |
| Kew | 268,886 | 109,493 |
| MdR | 360,954 | 110,351 |
| So | 326,742 | 226,241 |

**Supplementary Table S3.** Proportion of segregating sites per population [%].

| | *E. typhina* | *E. clarkii* |
|---|---|---|
| Aub | 55.0 | 51.9 |
| AubX | 48.2 | 47.7 |
| Auv | 55.9 | 46.3 |
| Cev | 55.0 | 47.6 |
| El | 53.9 | 42.1 |
| Kew | 40.9 | 27.6 |
| MdR | 54.9 | 27.8 |
| So | 49.7 | 56.9 |

A similar proportion of sites was polymorphic in most *E. clarkii* populations compared to *E. typhina* populations despite *E. clarkii* having a lower overall diversity. This could indicate similar mutation rates in the two species. Exceptions were Kew and MdR.

**Supplementary Table S4.** Fixed differences between populations.

| | | MdR | Kew | Auv | Cev | Aub | AubX | El | So | |
|---|---|---|---|---|---|---|---|---|---|---|
| *E. clarkii* | MdR | | 448 | 121 | 9 | 58 | 0 | 177 | 340 | *E. typhina* |
| | Kew | 10669 | | 22 | 137 | 26 | 197 | 24 | 690 | |
| | Auv | 4598 | 685 | | 0 | 0 | 83 | 0 | 214 | |
| | Cev | 5337 | 1922 | 83 | | 9 | 0 | 126 | 170 | |
| | Aub | 4827 | 1285 | 78 | 0 | | 0 | 0 | 11 | |
| | AubX | 5165 | 1493 | 58 | 23 | 0 | | 0 | 1 | |
| | El | 6214 | 1672 | 173 | 44 | 1 | 3 | | 57 | |
| | So | 5561 | 3507 | 762 | 21 | 288 | 263 | 486 | | |

Number of loci fixed for alternative alleles ($F_{ST}$=1) between population pairs of *E. typhina* (top right) and *E. clarkii* (bottom left). Populations are ordered according to longitude (West to East).

**Supplementary Table S5.** Mean nucleotide diversity ($\pi$) per site.

| | *E. typhina* | *E. clarkii* |
|---|---|---|
| Aub | 0.00269 | 0.00109 |
| AubX | 0.00265 | 0.00108 |
| Auv | 0.00249 | 0.00098 |
| Cev | 0.00258 | 0.00107 |
| El | 0.00256 | 0.00092 |
| Kew | 0.00240 | 0.00081 |
| MdR | 0.00249 | 0.00077 |
| So | 0.00241 | 0.00118 |

**Supplementary Table S6.** Mean Tajimas D calculated over 40kb windows.

| | *E. typhina* | *E. clarkii* |
|---|---|---|
| Aub | -0.334 | -0.211 |
| AubX | -0.366 | -0.354 |
| Auv | -0.369 | 0.102 |
| Cev | -0.061 | 0.151 |
| El | -0.307 | -0.015 |
| Kew | -0.093 | 0.903 |
| MdR | -0.183 | 1.318 |
| So | -0.282 | -0.076 |

**Supplementary Table S7.** Hybridization rates in two sympatric populations.

| | Dg_indv | Hl_indv | Et_collected | Ec_collected | total_collected | Et_probes | Ec_probes | total_probes | Et_hyb | Ec_hyb | Et_hyb_prop | Ec_hyb_prop | total_hyb | total_hyb_prop |
|---|---|---|---|---|---|---|---|---|---|---|---|---|---|---|
| Cev | 13 | 12 | 21 | 27 | 48 | 63 | 81 | 144 | 7 | 11 | 0.111 | 0.1358 | 18 | 0.1250 |
| Auv | 7 | 9 | 16 | 24 | 40 | 48 | 72 | 120 | 6 | 4 | 0.125 | 0.0555 | 10 | 0.0833 |

Columns specify the number of sampled individual plants [1,2], the number of stromata collected [3,4,5], the number of perithecia probes taken [6,7,8], the number of probes with hybrid asci [9,10], the proportion of hybrid asci [11,12], the total number of probes with hybrid asci [13] and the total hybrid proportion across both species [14]. Dg = *Dactylis glomerata*; Hl = *Holcus lanatus*; Et = E. typhina; Ec = *E. clarkii*

**Supplementary Table S8.** Individual plants assessed in Cev.

| plant_indv | stroma_samples | probes | hyb_asci |
|---|---|---|---|
| Dg1 | 2 | 6 | 0 |
| Dg2 | 1 | 3 | 0 |
| Dg3 | 2 | 6 | 2 |
| Dg4 | 2 | 6 | 1 |
| Dg5 | 3 | 9 | 0 |
| Dg6 | 1 | 3 | 0 |
| Dg7 | 1 | 3 | 0 |
| Dg8 | 2 | 6 | 3 |
| Dg9 | 1 | 3 | 0 |
| Dg10 | 1 | 3 | 1 |
| Dg11 | 2 | 6 | 0 |
| Dg12 | 2 | 6 | 0 |
| Dg13 | 1 | 3 | 0 |
| Hl1 | 2 | 6 | 0 |
| Hl2 | 2 | 6 | 2 |
| Hl3 | 3 | 9 | 3 |
| Hl4 | 3 | 9 | 0 |
| Hl5 | 2 | 6 | 0 |
| Hl6 | 1 | 3 | 0 |
| Hl7 | 3 | 9 | 0 |
| Hl8 | 2 | 6 | 0 |
| Hl9 | 3 | 9 | 4 |
| Hl10 | 2 | 6 | 0 |
| Hl11 | 1 | 3 | 0 |
| Hl12 | 3 | 9 | 2 |
| Total | 48 | 144 | 18 |

**Supplementary Table S9.** Individual plants assessed in Auv.

| plant_indv | stroma_samples | probes | hyb_asci |
|---|---|---|---|
| Dg1 | 3 | 9 | 1 |
| Dg2 | 3 | 9 | 0 |
| Dg3 | 2 | 6 | 0 |
| Dg4 | 2 | 6 | 0 |
| Dg5 | 3 | 9 | 2 |
| Dg6 | 2 | 6 | 3 |
| Dg7 | 1 | 3 | 0 |
| Hl1 | 3 | 9 | 0 |
| Hl2 | 3 | 9 | 3 |
| Hl3 | 3 | 9 | 0 |
| Hl4 | 3 | 9 | 0 |
| Hl5 | 2 | 6 | 0 |
| Hl6 | 2 | 6 | 0 |
| Hl7 | 3 | 9 | 1 |
| Hl8 | 2 | 6 | 0 |
| Hl9 | 3 | 9 | 0 |
| Total | 40 | 120 | 10 |

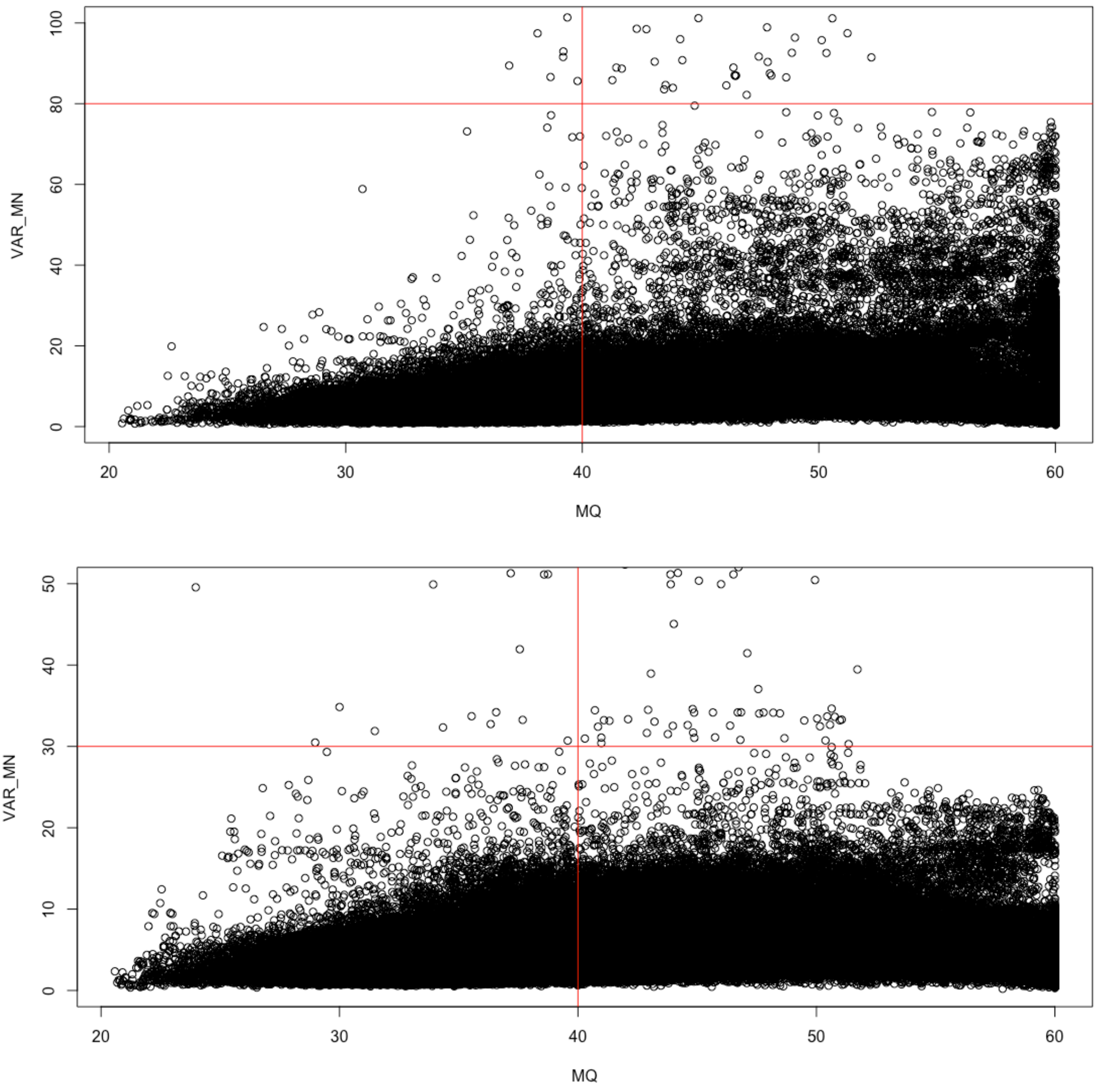


**Supplementary Figure S1.** Defining filtering cutoffs. We retained SNPs with a MQ value > 40 (the Root Mean Square of the mapping quality of the reads across all samples), and with a relative variance in read depth < 80 in *E. typhina* (top) and >30 in *E. clarkii* (bottom).

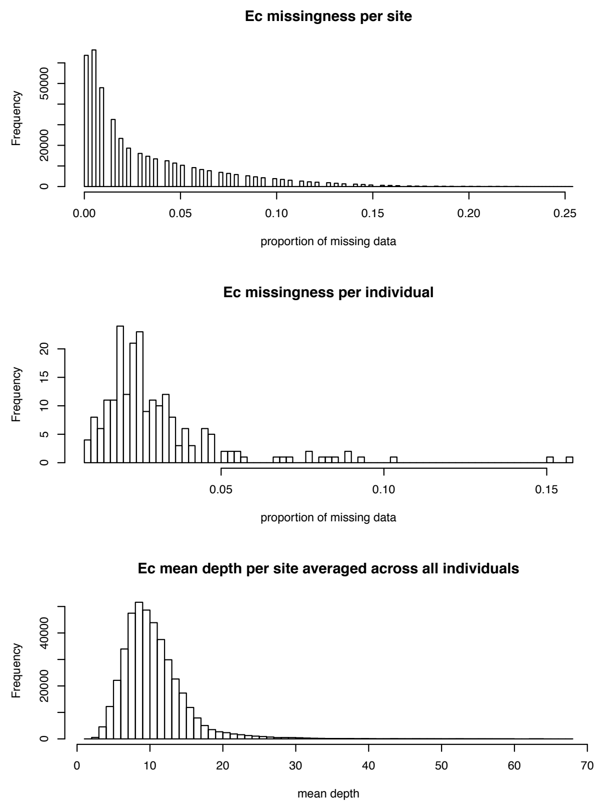


**Supplementary Figure S2.** Summary of statistics after filtering. For the *E. typhina* (right) and *E. clarkii* (left) SNP datasets, missingness per site, missingness per individual and mean depth per site are plotted as histograms. *E. clarkii* has more missing data and lower depth which is probably related to the slightly lower coverage with the larger reference genome and larger proportion of polymorphic regions where accurate mapping is more challenging.

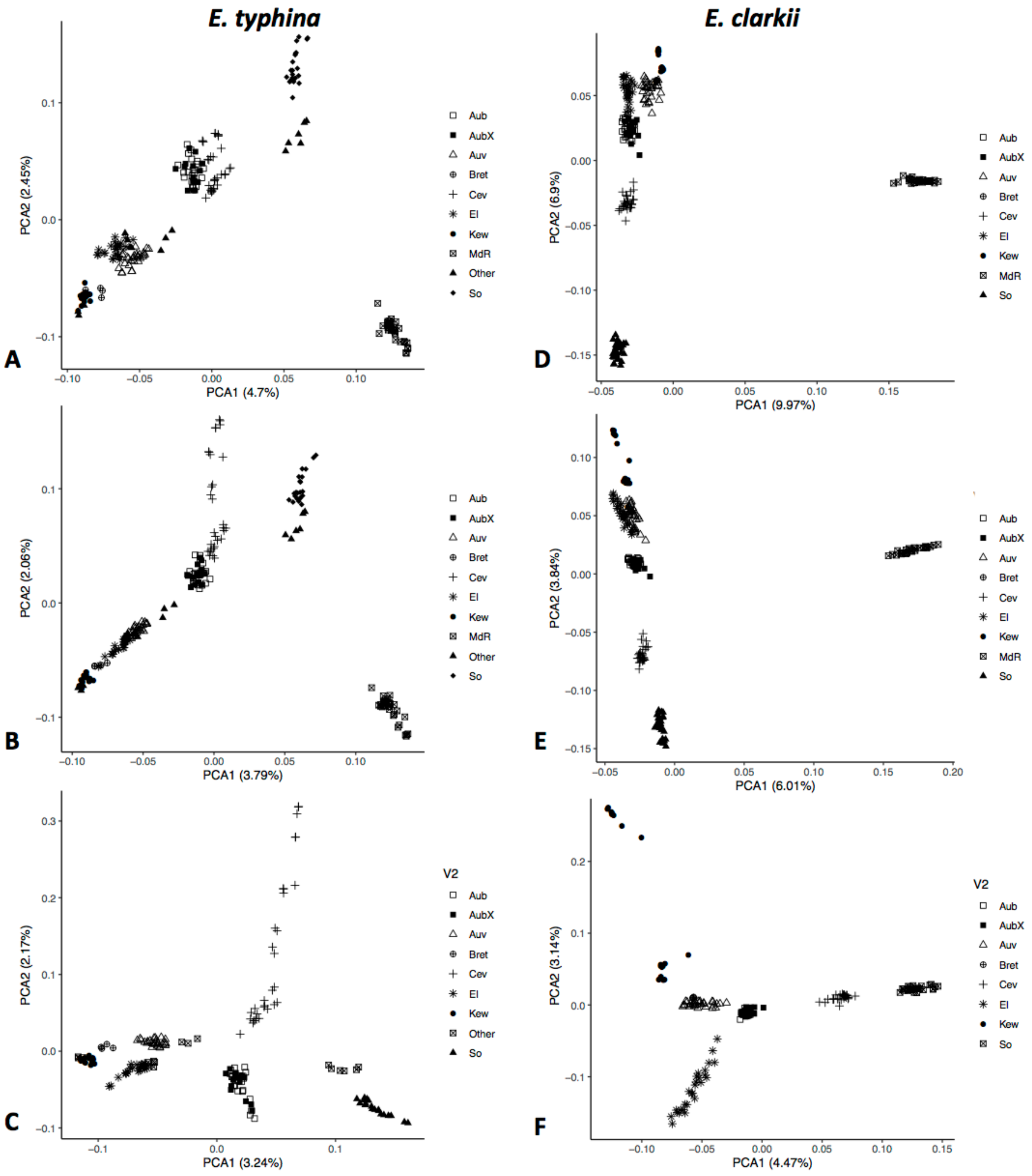


**Supplementary Figure S3.** PCAs based on SNP datasets with different filtering criteria. Percentage of variance explained by the first two principal components is shown in parentheses; symbols indicate sampling locations of populations. A, D. complete datasets including Spanish populations (MdR). B, E. LD pruned data with maf ≥ 0.05 filter. C, F. Same as B and E but with MdR removed.

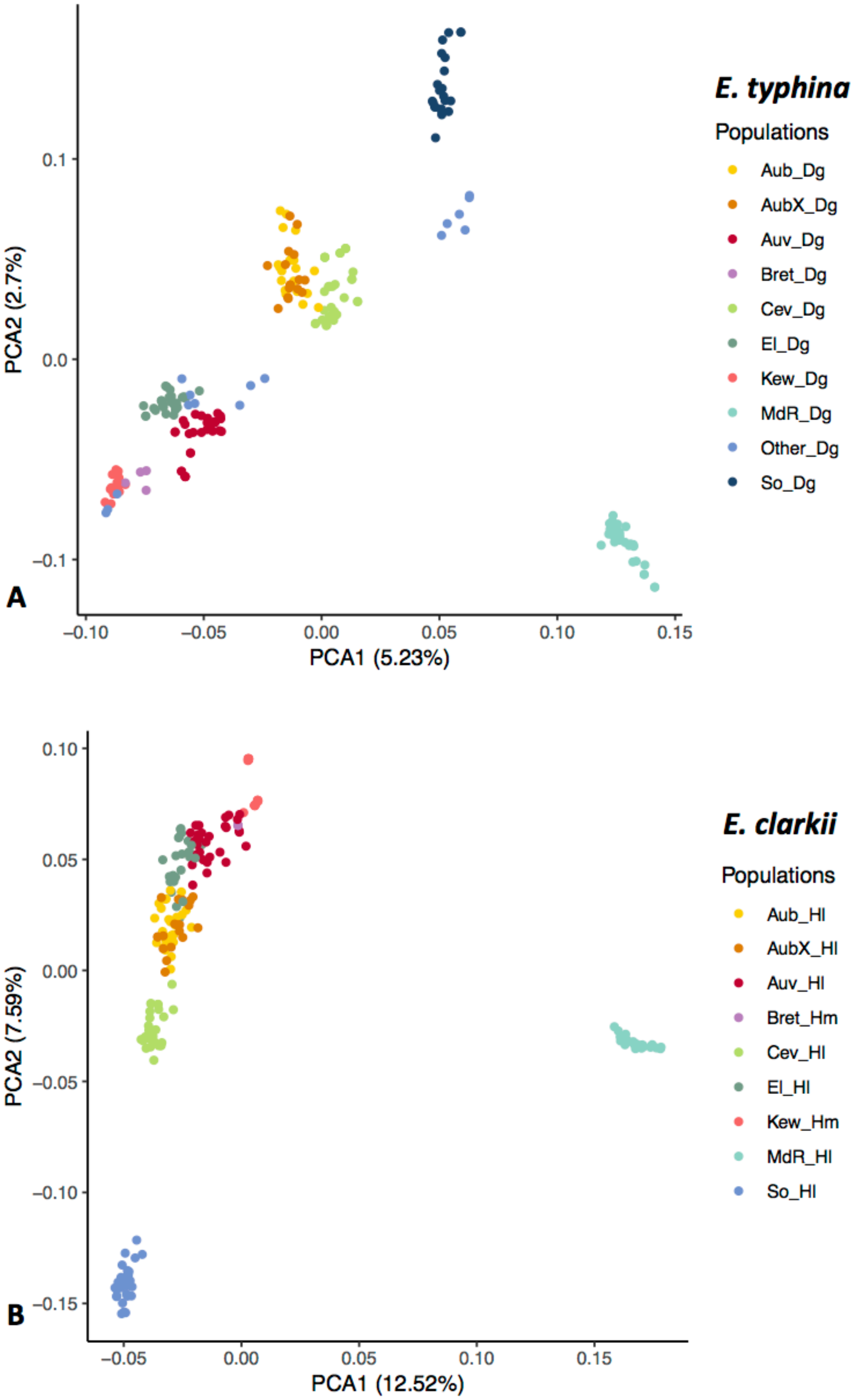


**Supplementary Figure S4.** PCAs based on SNP datasets with no missing data. Percentage of variance explained by the first two principal components is shown in parentheses; colors indicate sampling locations of populations.

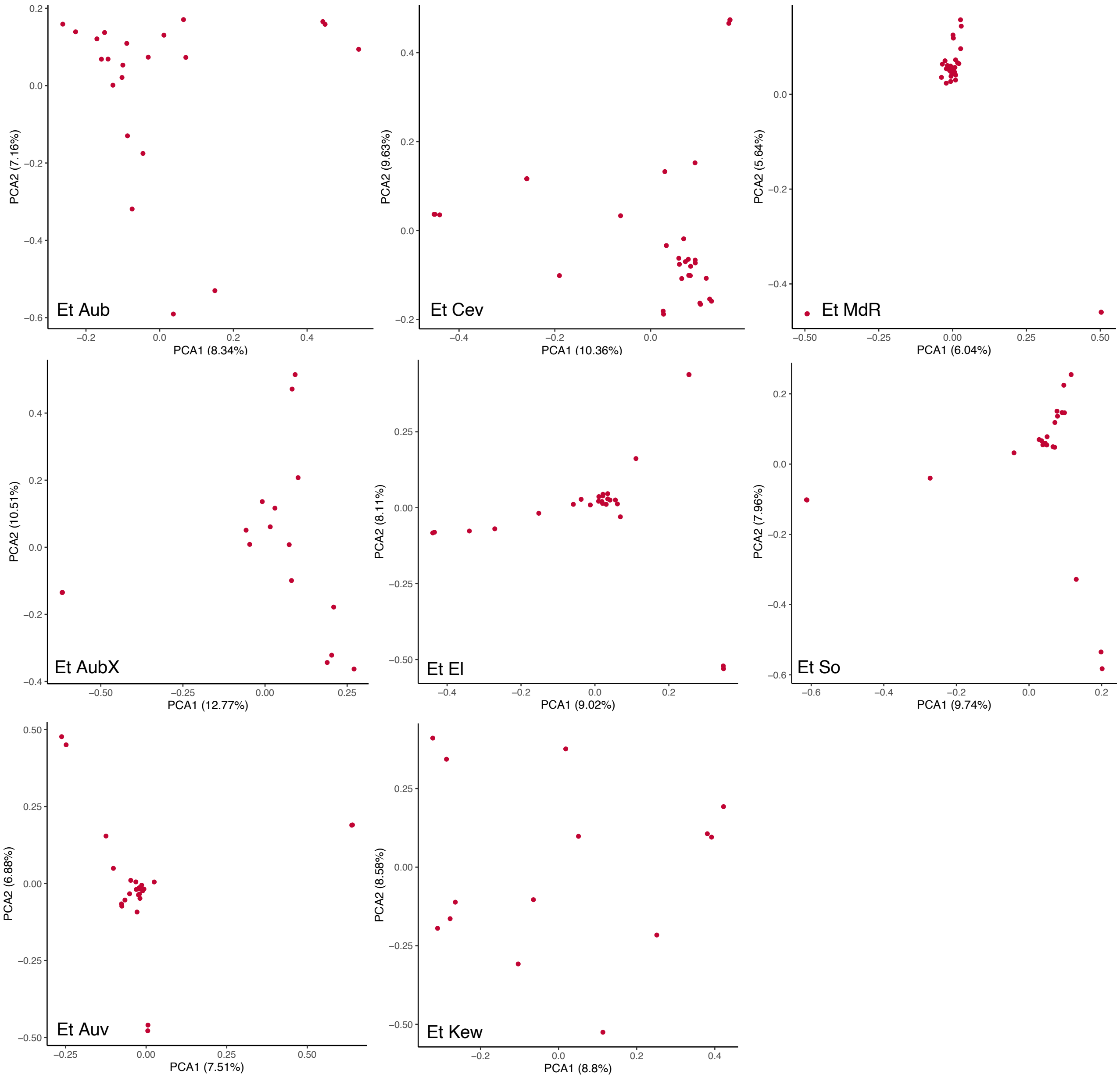


**Supplementary Figure S5.** PCAs within populations of *E. typhina*.

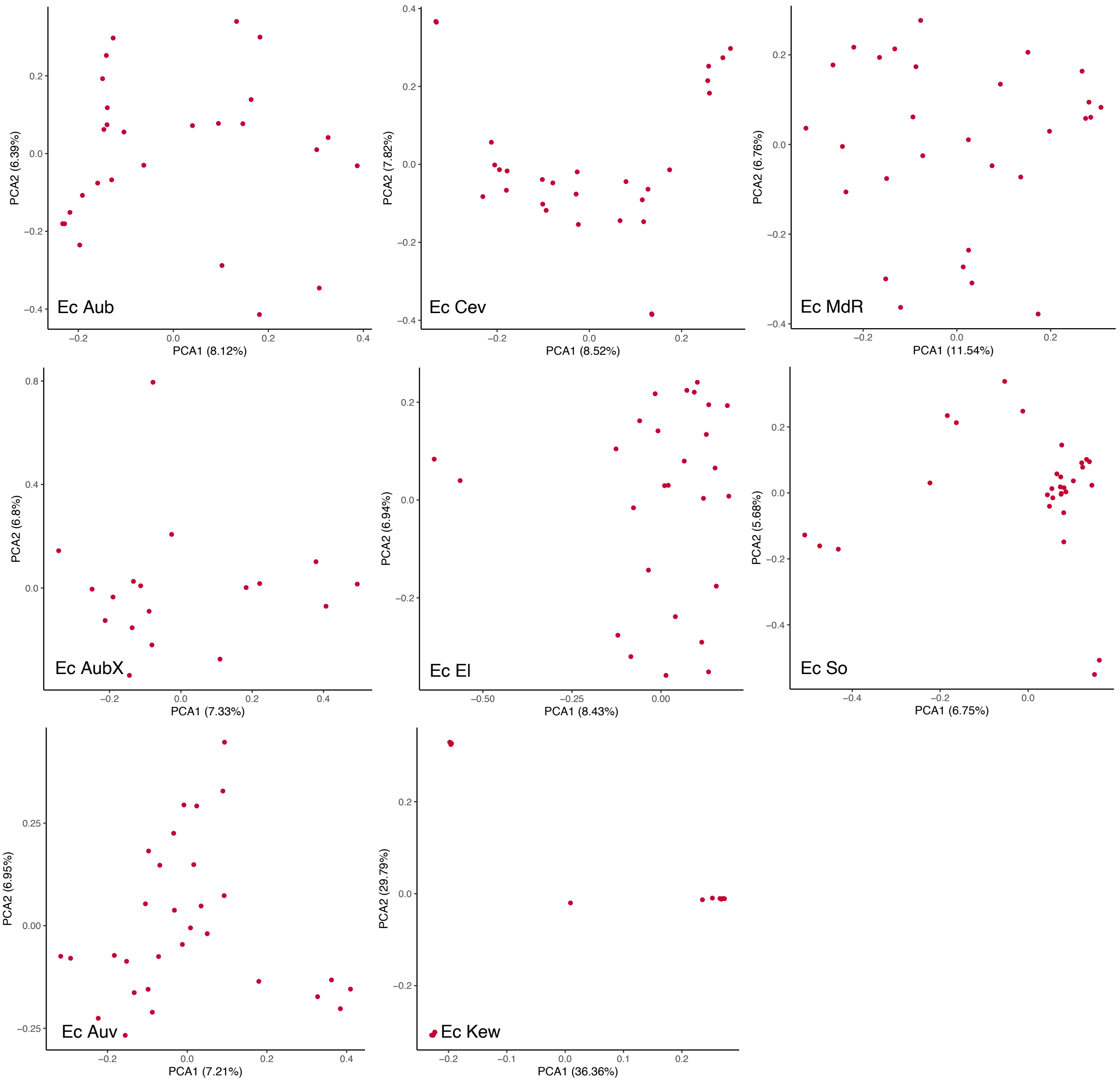


**Supplementary Figure S6.** PCAs within populations of *E. clarkii*.

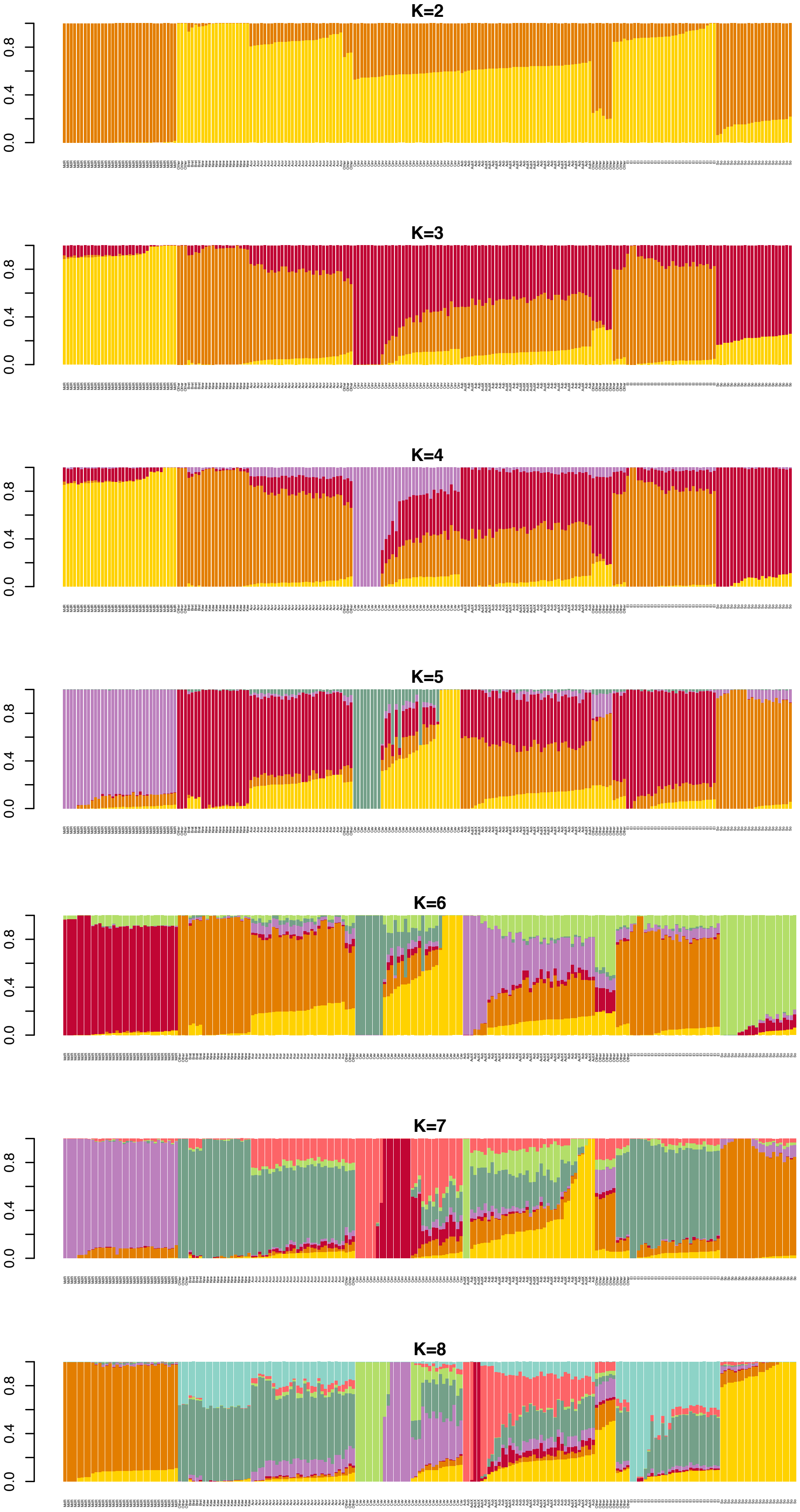
K=2
0.0
0.4
0.8
K=3
0.0
0.4
0.8
K=4
0.0
0.4
0.8
K=5
0.0
0.4
0.8
K=6
0.0
0.4
0.8
K=7
0.0
0.4
0.8
K=8
0.0
0.4
0.8

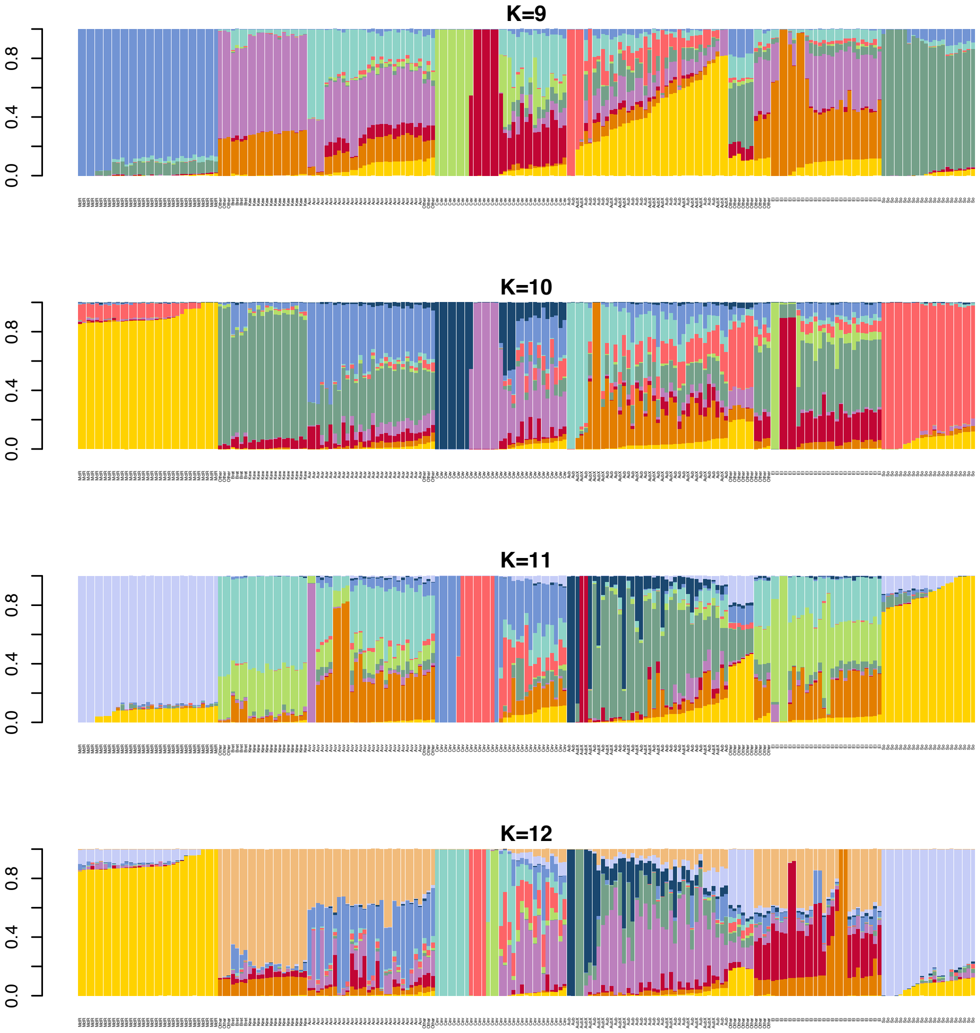


**Supplementary Figure S7.** *E. typhina* STRUCTURE results for all Ks. Bar plots of membership proportions (y-axis) as inferred by STRUCTURE for K=2 – K=12.

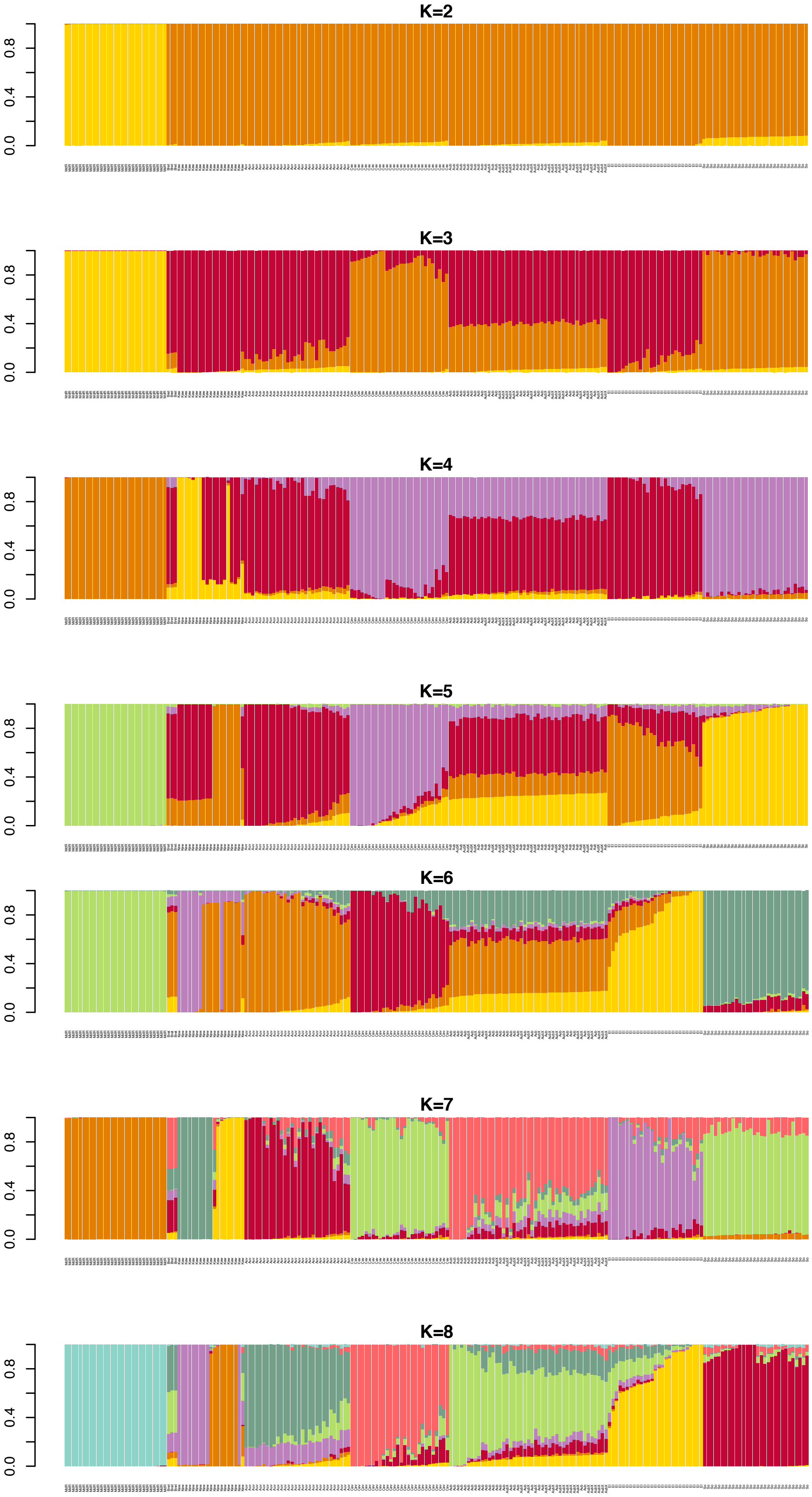
K=2
K=3
K=4
K=5
K=6
K=7
K=8
0.0
0.4
0.8

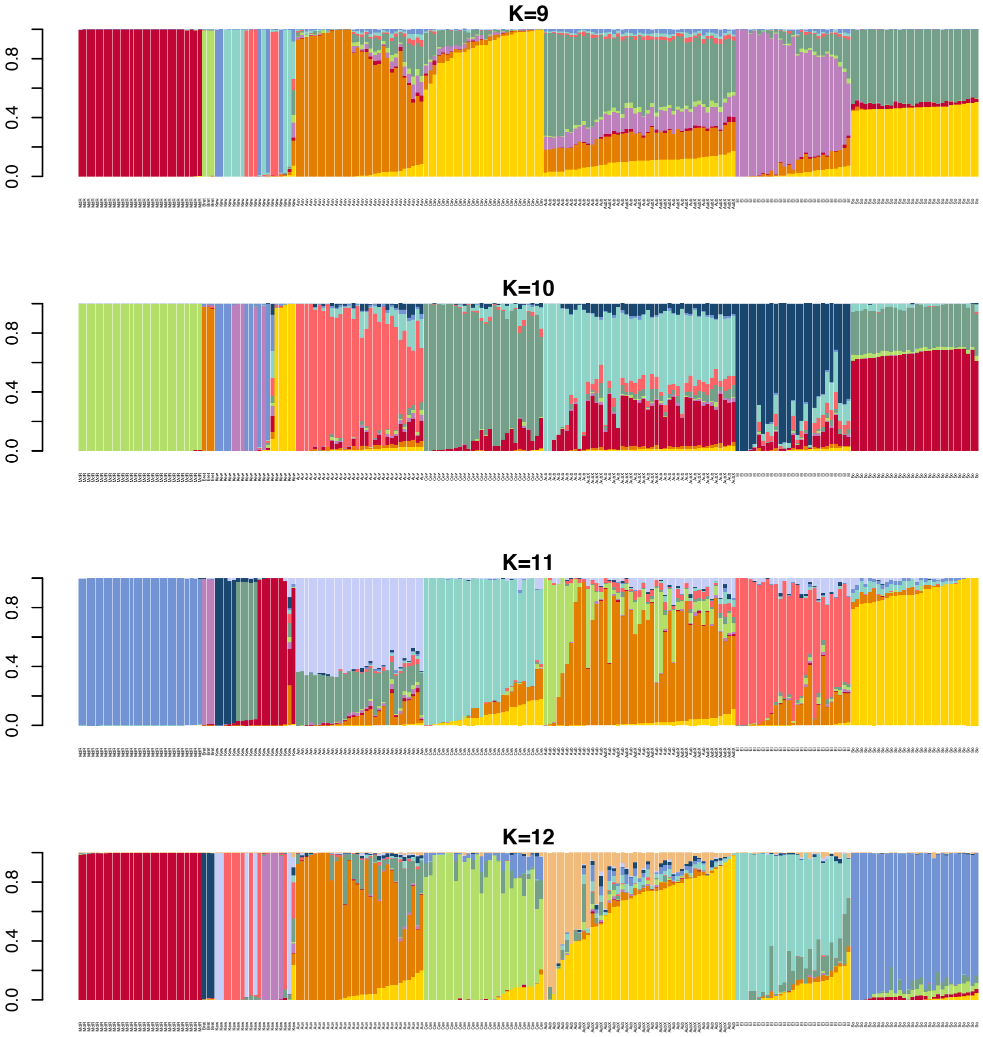


**Supplementary Figure S8.** *E. clarkii* STRUCTURE results for all Ks. Bar plots of membership proportions (y-axis) as inferred by STRUCTURE for K=2 – K=12.

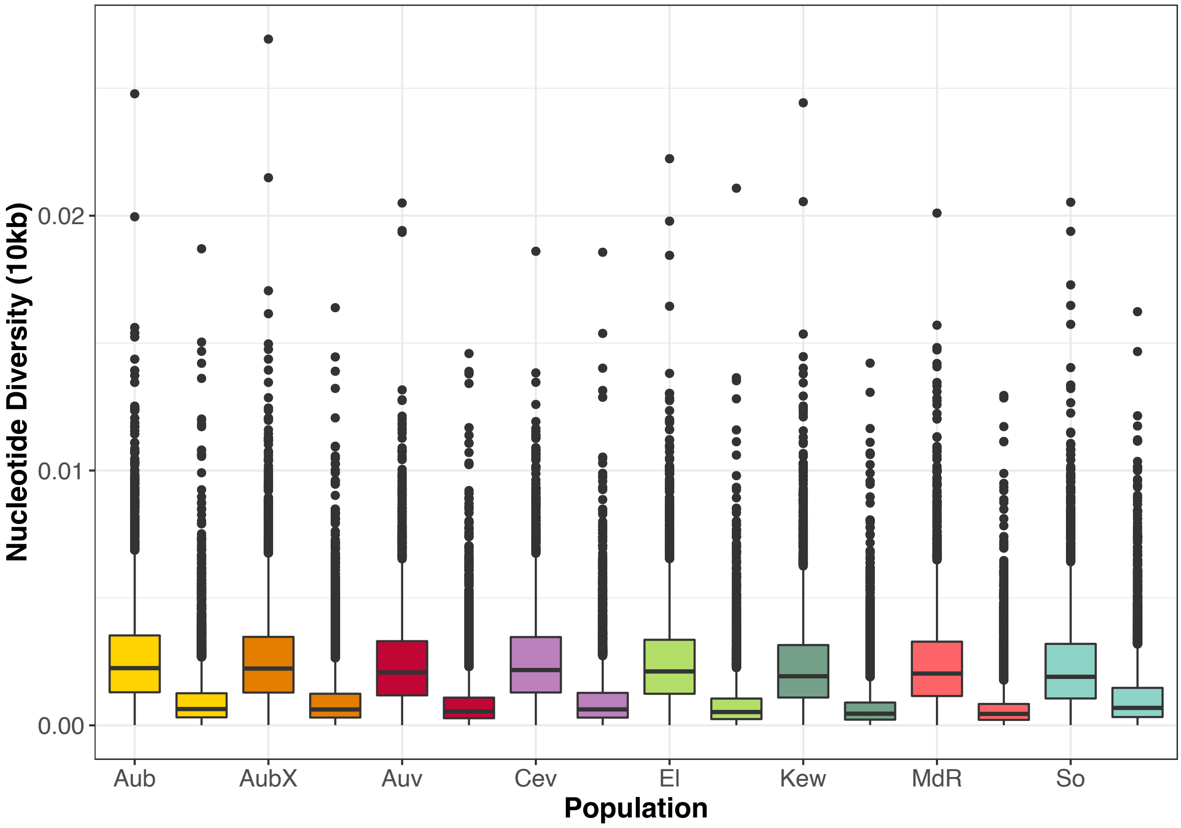


**Figure S9.** Nucleotide diversity per population. Nucleotide diversity (π) calculated in 10kb sliding windows along genomes. Colors correspond to sympatric population pairs, with the first boxplot of each color showing *E. typhina* and the second *E. clarkii*. Lower and upper hinges correspond to the first and third quartiles (the 25th and 75th percentiles); whiskers extend to 1.5 times the inter-quartile range and outlier data beyond this value are plotted as individual points.

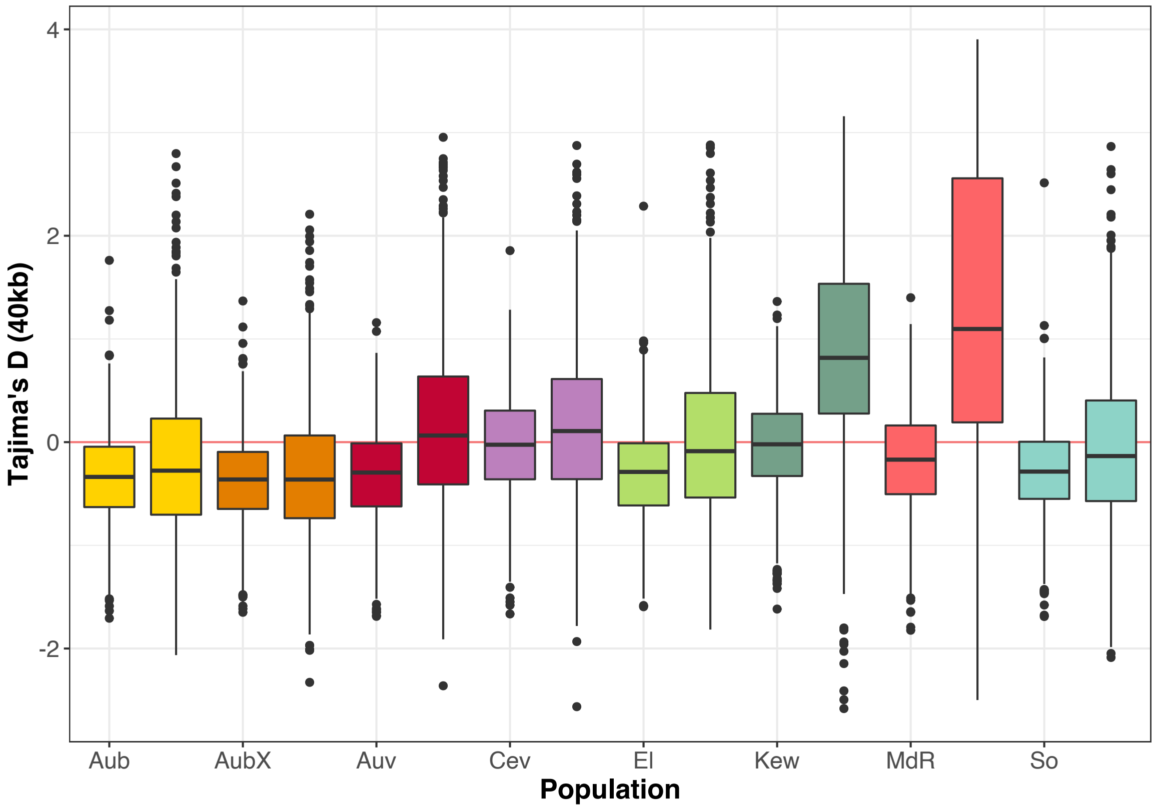


**Supplementary Figure S10.** Tajima's D per population. Tajima's D calculated in 40kb sliding windows along genomes. Colors correspond to sympatric population pairs, with the first boxplot of each color showing *E. typhina* and the second *E. clarkii*. Lower and upper hinges correspond to the first and third quartiles (the 25th and 75th percentiles); whiskers extend to 1.5 times the inter-quartile range and outlier data beyond this value are plotted as individual points.

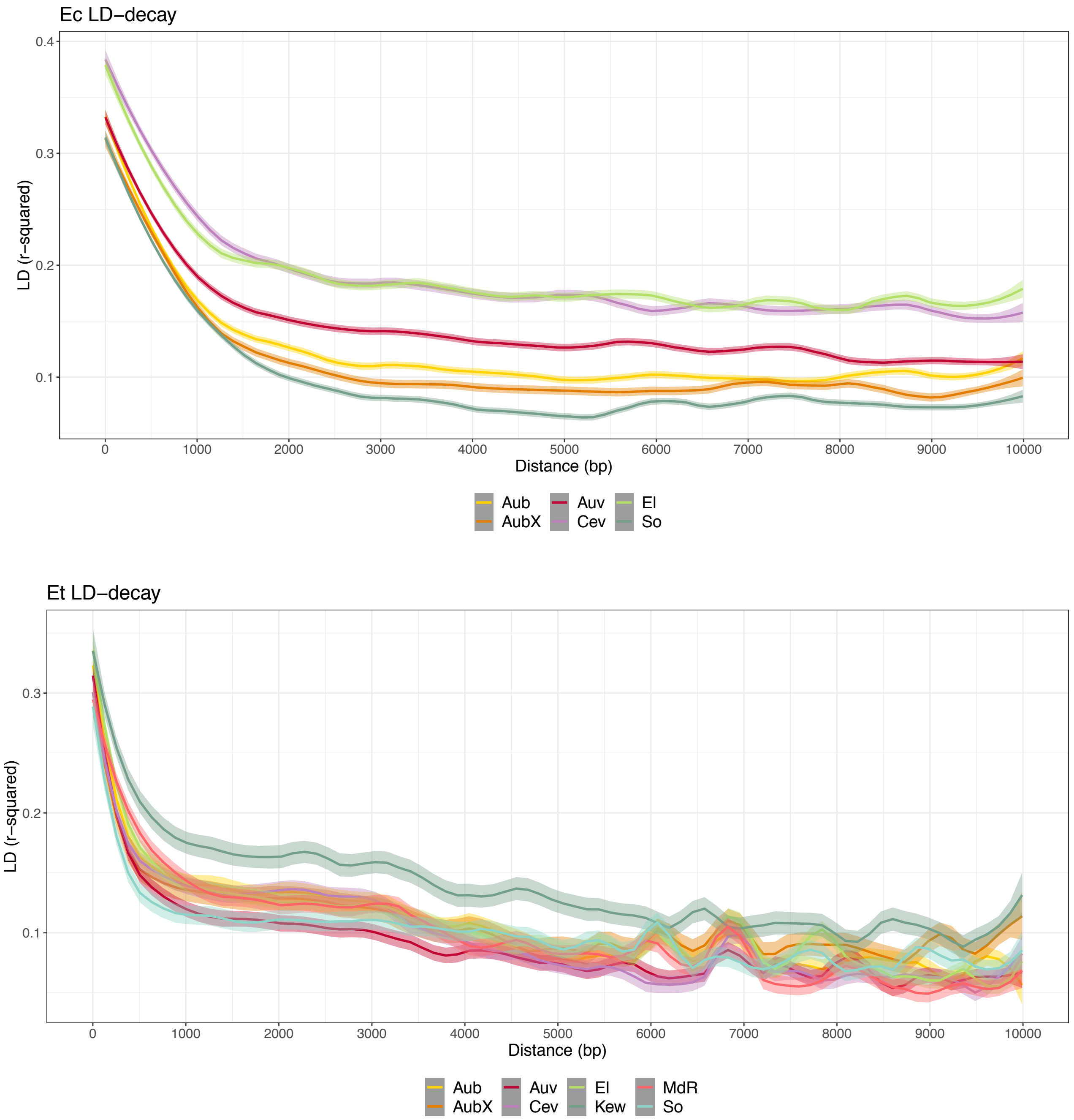


**Supplementary Figure S11.** Decay of linkage disequilibrium ($r^2$) with physical distance in *E. typhina* (top) and *E. clarkii* (bottom) populations.

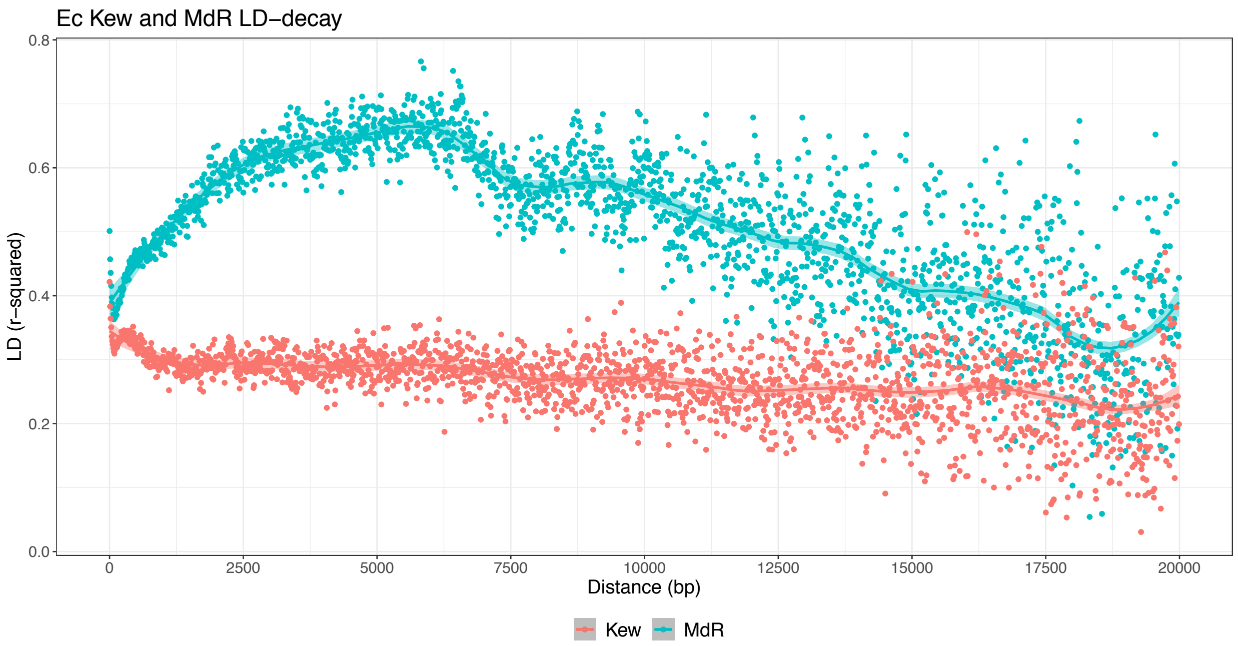


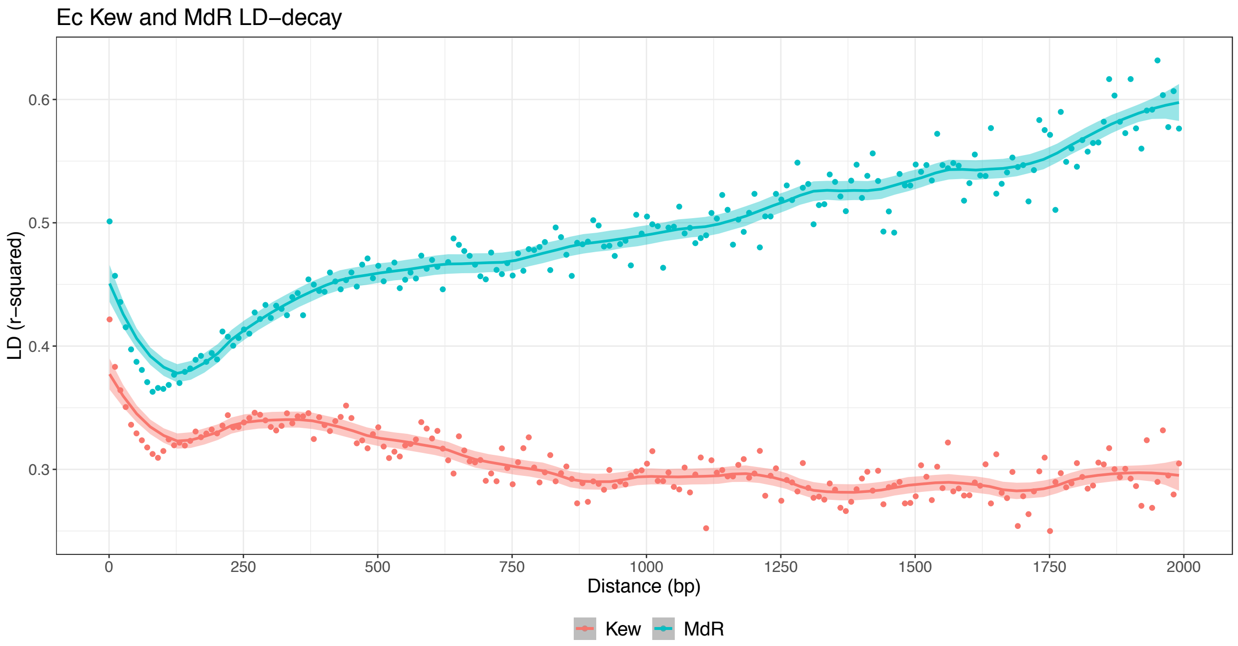


**Supplementary Figure S12.** Decay of linkage disequilibrium ($r^2$) with physical distance in *E. clarkii* Kew and MdR. LD decayed very slowly in Kew (red) and MdR had high LD overall. The bottom plot is zoomed in on the first 2kb showing an initial LD decay within the first 100bp.

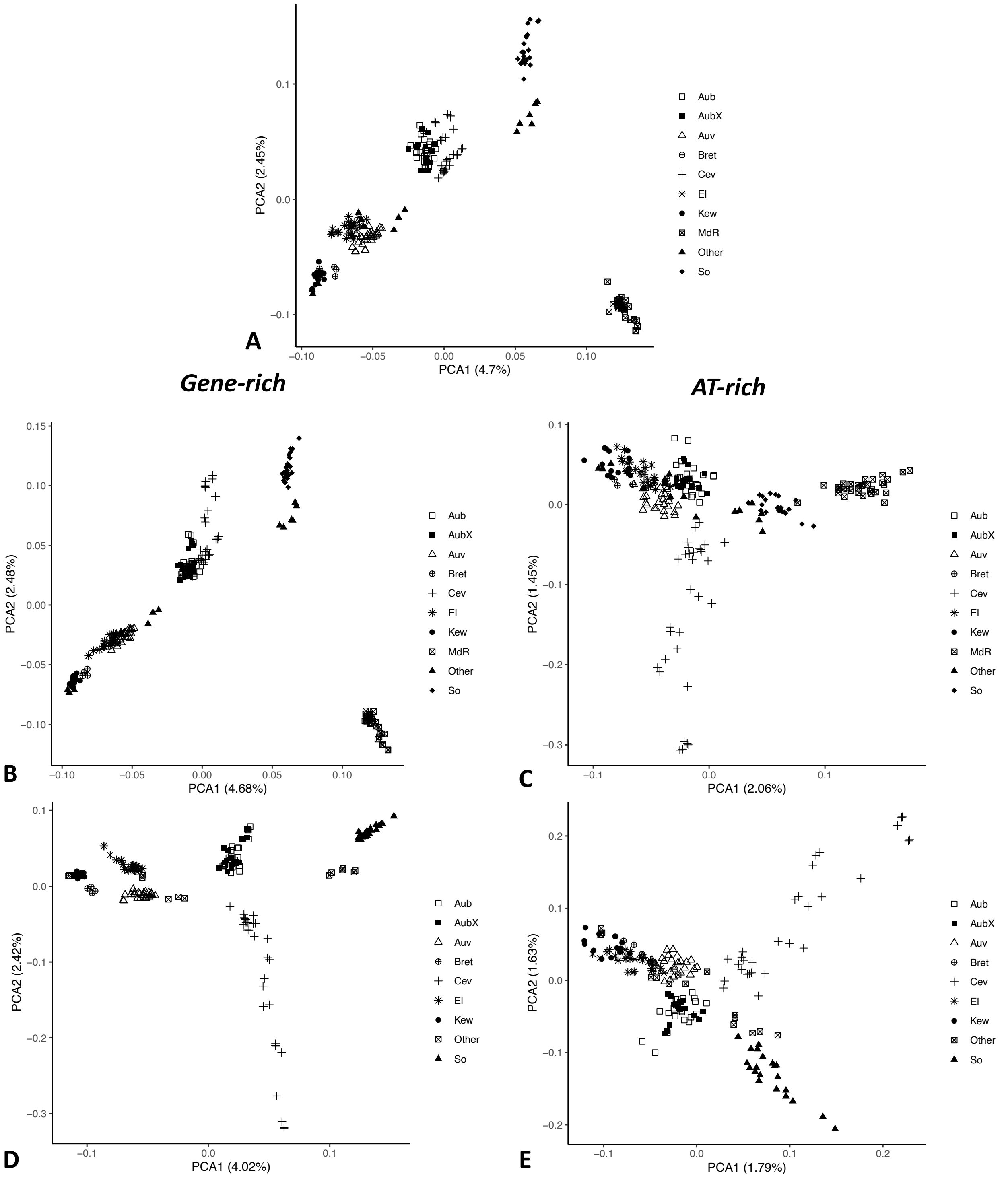


**Supplementary Figure S13.** PCAs by genome compartments in *E. typhina*. Analyses were performed with complete SNP datasets split by compartments. For reference the PCA for complete genome-wide SNPs including MdR is shown (**A**, same as Supplementary Figure S3A). **(B, C)** Compartment PCAs including MdR. **(D, E)** Compartment PCAs excluding MdR. Percentage of variance explained by the first two principal components is shown in parentheses; symbols indicate sampling locations of populations.

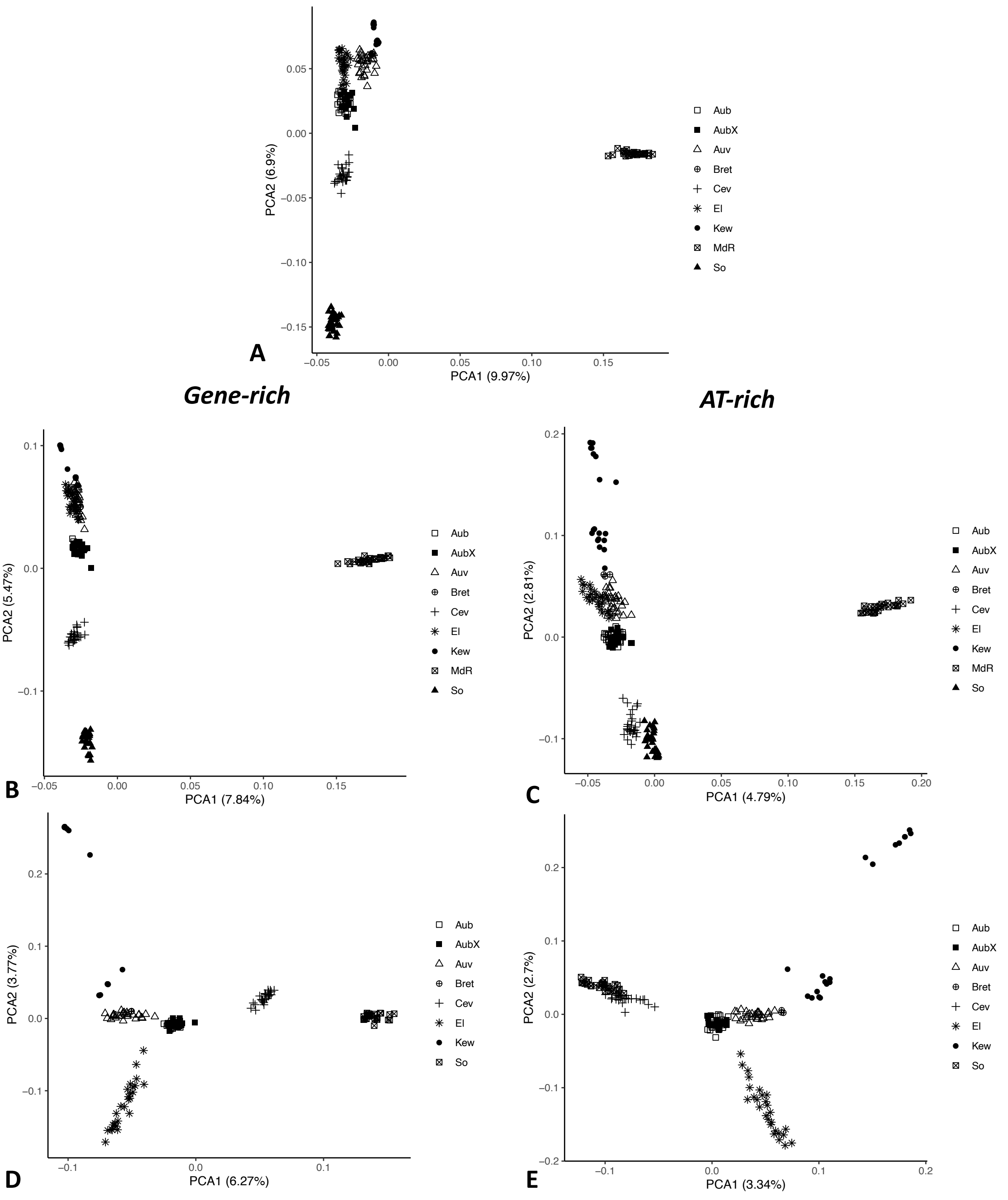


**Supplementary Figure S14.** PCAs by genome compartments in *E. clarkii*. Analyses were performed with complete SNP datasets split by compartments. For reference the PCA for complete genome-wide SNPs including MdR is shown (**A**, same as Supplementary Figure S3D). **(B, C)** Compartment PCAs including MdR. **(D, E)** Compartment PCAs excluding MdR. Percentage of variance explained by the first two principal components is shown in parentheses; symbols indicate sampling locations of populations.

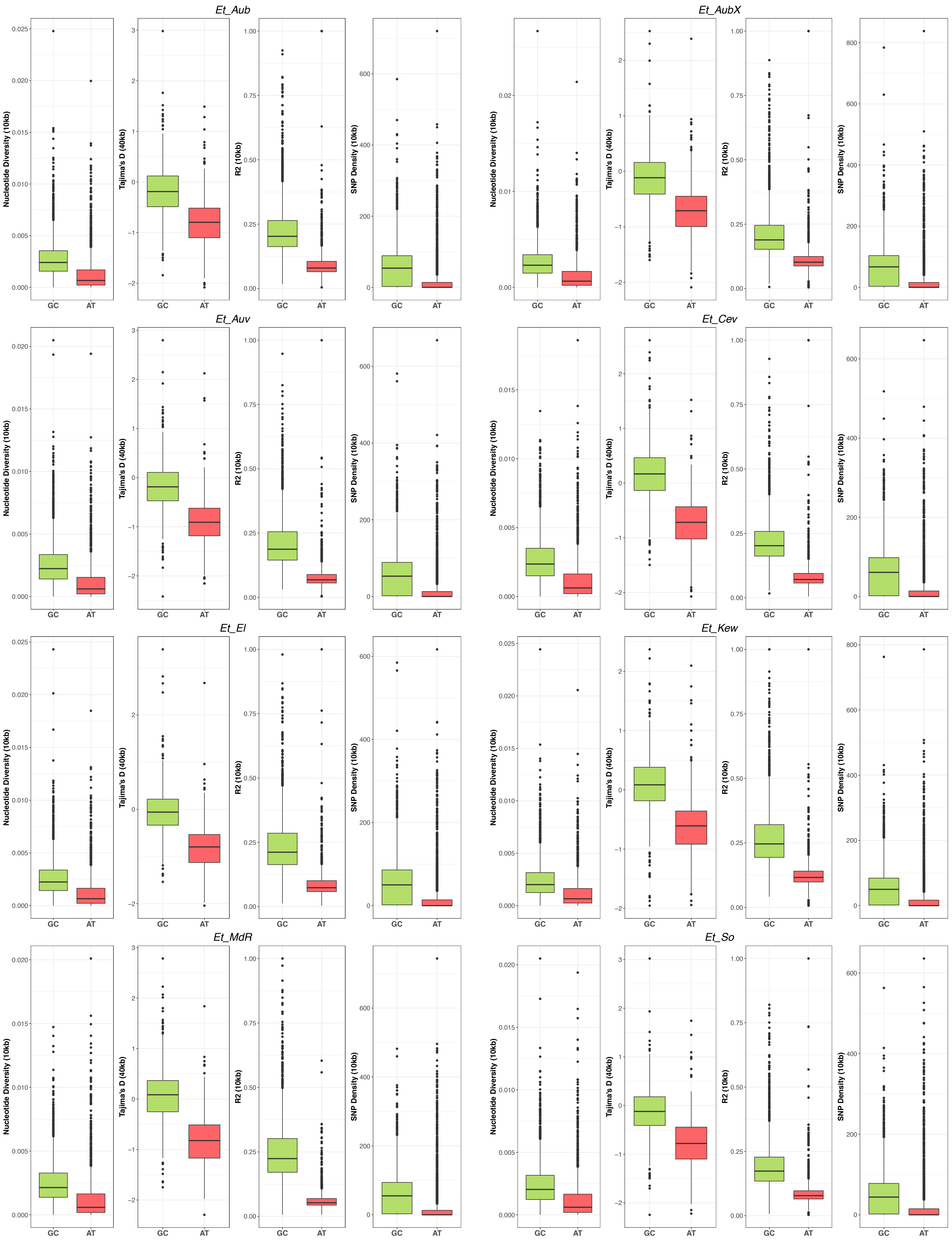

Et_Aub
Et_AubX
Et_Auv
Et_Cev
Et_El
Et_Kew
Et_MdR
Et_So
Nucleotide Diversity (10kb)
Tajima's D (40kb)
R2 (10kb)
SNP Density (10kb)
GC
AT

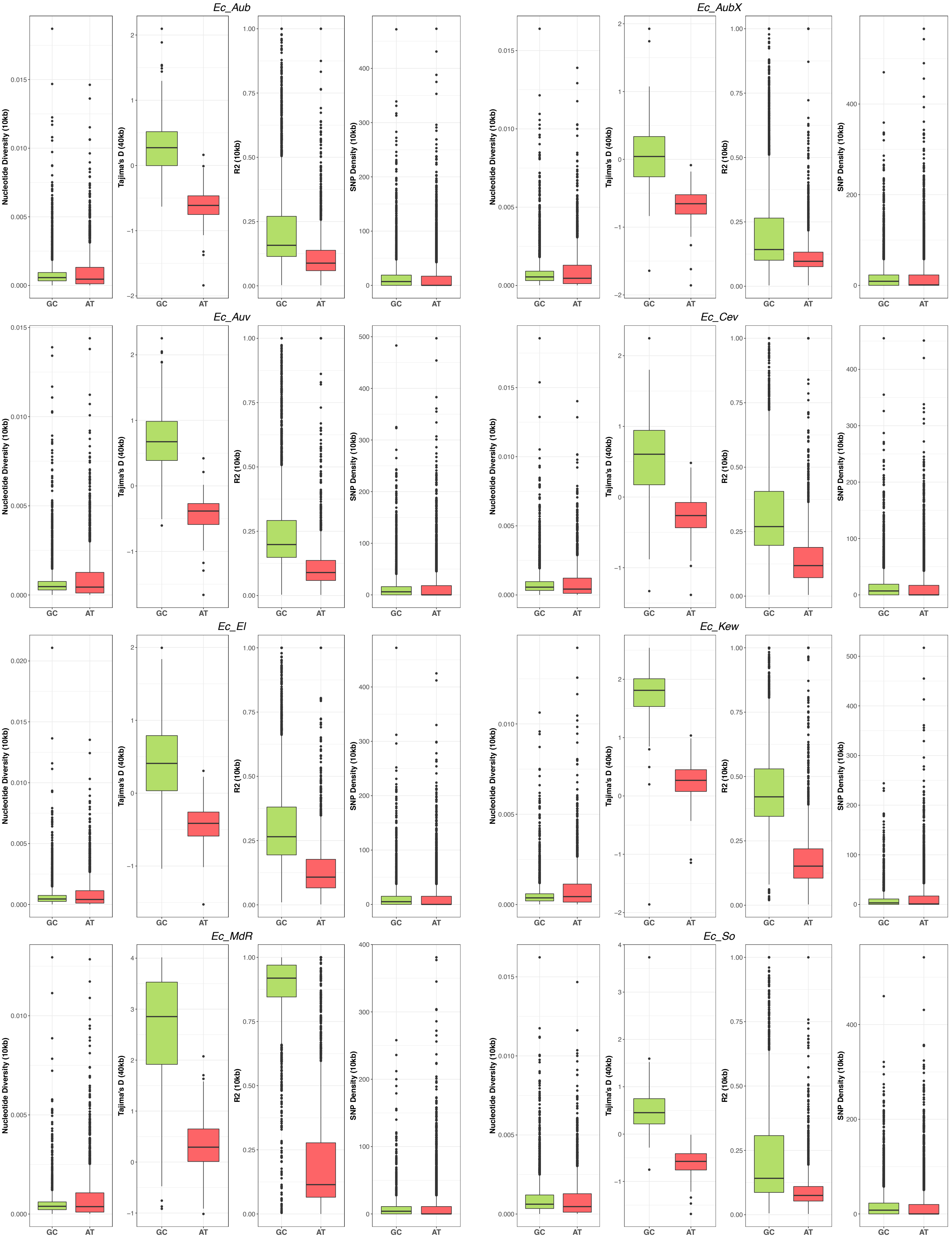

Ec_Aub
Ec_AubX
Ec_Auv
Ec_Cev
Ec_El
Ec_Kew
Ec_MdR
Ec_So
Nucleotide Diversity (10kb)
Tajima's D (40kb)
R2 (10kb)
SNP Density (10kb)
GC
AT

Previous two pages:

**Supplementary Figures S15 and S16.** Population summary statistics by genome compartments. In each panel statistics for one population are shown as boxplots, green for the gene-rich compartment (GC) and red for the AT-rich compartment (AT). From left to right: Nucleotide diversity ($\pi$) in 10kb windows, Tajima's D in 40kb windows, LD $r^2$ in 10kb windows, SNP density in 10kb windows Species and population IDs are indicated in the headers (Et = *E. typhina* and Ec = *E. clarkii*). Lower and upper hinges correspond to the first and third quartiles (the 25th and 75th percentiles); whiskers extend to 1.5 times the inter-quartile range and outlier data beyond this value are plotted as individual points.

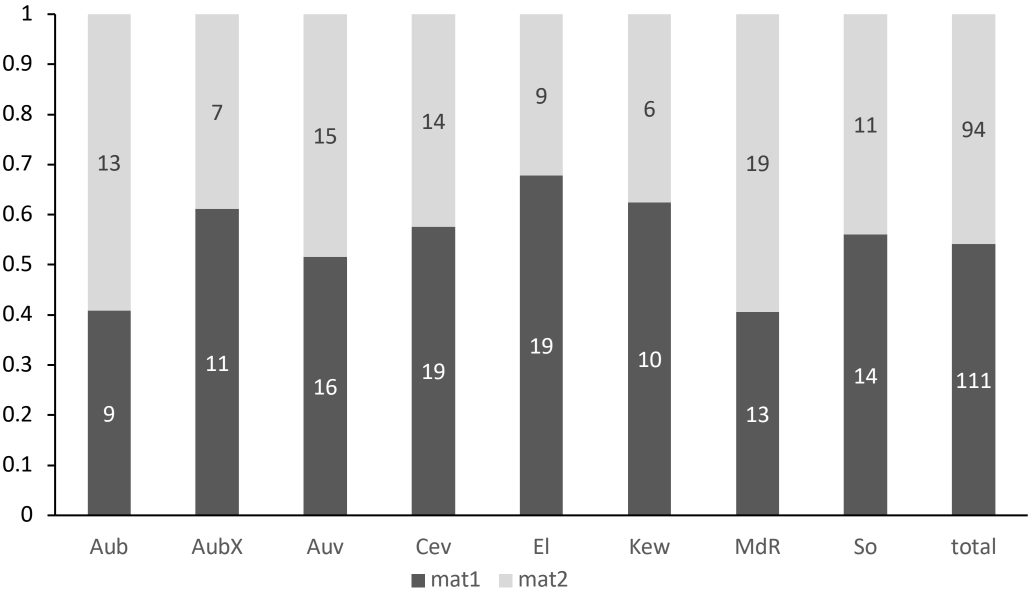


**Supplementary Figure S17.** Proportion of mating types in *E. typhina* populations and across all sequenced genotypes. Numbers indicate how many individuals had mat-1 (dark grey) and how many had mat-2 (light grey).

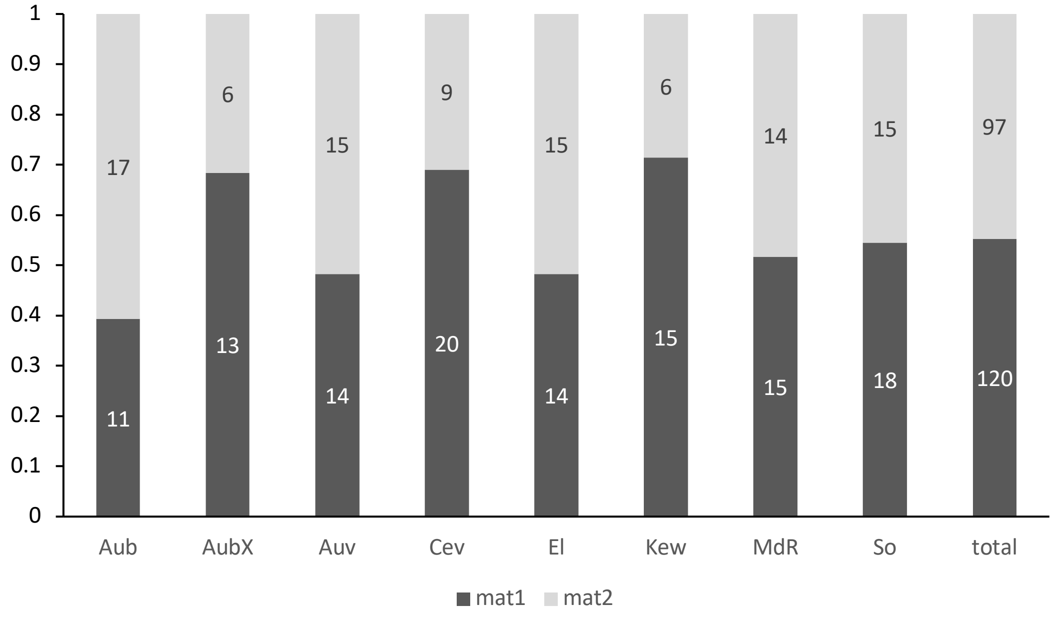


**Supplementary Figure S18.** Proportion of mating types in *E. clarkii* populations and across all sequenced genotypes. Numbers indicate how many individuals had mat-1 (dark grey) and how many had mat-2 (light grey).
